# Systematic Approach Identifies Multiple Transcription Factor Perturbations That Rejuvenate Replicatively Aged Human Skin Fibroblasts

**DOI:** 10.1101/2022.11.20.517270

**Authors:** Janine Sengstack, Jiashun Zheng, Michael Mobaraki, Jue Lin, Changhui Deng, Hao Li

## Abstract

Rejuvenation, long a quixotic dream, recently became a possibility through exciting new approaches to counteract aging. For example, parabiosis and partial reprogramming through overexpressing four stem cell transcription factors (Yamanaka factors) both rejuvenate organisms and cells^1–5^. We hypothesize there are many other genetic solutions to human cell rejuvenation, and some solutions may be safer and more potent than current gene targets. We set out to develop a systematic approach to identify novel genes that, when overexpressed or repressed, reprogram the global gene expression of a cell back to a younger state. Using the Hayflick model of human cell replicative aging, we performed a Perturb-seq screen of 200 transcription factors (TFs) selected through a combination of bioinformatic analysis and literature search. We identified dozens of potentially rejuvenating TFs—those that when overexpressed or repressed in late passage cells reprogrammed global gene expression patterns back to an earlier passage state. We further validated four top TF perturbations through molecular phenotyping of various aging hallmarks. Late passage cells either overexpressing EZH2 or E2F3 or repressing STAT3 or ZFX had more cell division, less senescence, improved proteostasis, and enhanced mitochondrial function. These TF perturbations led to similar downstream gene expression programs. In addition, the rejuvenating effects of these TFs were independent of telomeres. We believe our general approach for identifying rejuvenating factors can be applied to other model systems, and some of the top TF perturbations we discovered will lead to future research in novel, safer rejuvenation therapies.

## Introduction

The idea of rejuvenation to counteract aging is as old as human civilization. For millennia, humankind has dreamt of rejuvenation therapies that could reverse aging, and tales of the fountain of youth have been recounted across continents. But, without a cellular level understanding of aging coupled with advanced intervention technologies, rejuvenation remained a fiction. Recent breakthroughs in aging research changed this situation drastically and brought us the hope that rejuvenation may become a reality. For example, systemic factors in young blood rejuvenate various mouse tissues and brain function^1,2,6^, and partial reprogramming with four stem cell transcription factors (OCT4, SOX2, KLF4, and MYC, or the Yamanaka factors) rejuvenates both tissues and cells and extends the lifespan of mice^3–5^. More recently, it was found that cyclic induction of an N terminal truncated form of FOXM1 in mice delays natural and progeroid aging phenotypes and extends their lifespan^7^.

These discoveries support the notion that “young” and “old” can be described as different gene expression states, and the “old” state can be reversed back into a “young” state. The Yamanaka factors, arguably the most famous and well-studied rejuvenation factors^3,4^, are being intensely studied by academics and biotechnology companies alike^8^. Unfortunately, there is considerable cancer risk when overexpressing the Yamanaka factors. Previous work suggests cancer risk can be minimized by optimizing the dose and schedule of gene induction^3,5,7^. While an exciting proof of concept, it is very challenging to translate this rejuvenation protocol to humans.

We hypothesized that there are many other genetic solutions to human cell rejuvenation, and some solutions may be safer and more potent than current gene targets. Finding more genetic solutions is also important as it will increase the chance of successful future translation into human therapy. We set out to develop a systematic approach to identify novel genes that, when overexpressed or repressed, reprogram the global gene expression state of a cell back to a younger state. We decided to focus on transcription factors and chromatin modifiers because they influence the expression of many other genes (hereafter we will call them TFs for the ease of description).

To test our hypothesis, we utilized a canonical model for human cell aging and senescence, passaged human skin fibroblasts. Leonard Hayflick discovered that fibroblast cells gradually age *in vitro*, eventually becoming senescent after about 40 to 60 population doublings (PD)—a phenomenon termed the Hayflick limit^9^. We used early, middle, and late passage fibroblasts to model young, middle-aged, and old cells, respectively, because these passaged cells display aspects of both cellular aging and senescence^10–13^.

We first performed single-cell RNA-sequencing (scRNA-seq) on passaged fibroblasts to define their gene expression states. Next, we aimed to find TF perturbations that could change the global gene expression of a late passage cell back to an earlier passage state. Using a combination of bioinformatic analyses of differentially expressed TF modules and literature searches, we selected 200 candidate TFs. We next performed a high-throughput Perturb-seq^14^ screen of these TFs in late passage cells, overexpressing and repressing each TF individually with CRISPRa (CRA)^14^ and CRISPRi (CRI)^15^ respectively, and measured gene expression changes via scRNA-seq. We reasoned that using global gene expression as a high-dimensional readout would give us a higher likelihood of success than using one or two gene reporters, as cell aging is a complex phenotype involving scores of genes. Amazingly, over a dozen TF perturbations reversed global gene expression in late passage cells back to an earlier state. Further TF module analyses^16^ indicated these top TF perturbations caused similar, convergent gene expression changes, even though the TFs themselves originated from diverse upstream pathways.

We further tested our top TF perturbations from the Perturb-seq screen with comprehensive cell and molecular phenotyping, and we found four TF perturbations which reversed various cell aging hallmarks^11^. Late passage CRA cells overexpressing EZH2 or E2F3, and late passage CRI cells repressing STAT3 or ZFX had more cell division, less senescence, improved proteostasis, and enhanced mitochondrial function. No combination or cocktail of gene perturbations was required, and the fibroblasts always maintained their cell identity. In addition, the rejuvenating effects of these TFs were independent of telomeres.

We believe our general approach for identifying rejuvenating factors can be applied to other model systems, and some of the top TF perturbations we discovered will lead to future research in novel, safer rejuvenation therapies.

## Results

### Characterizing the gene expression states of passaged human fibroblast cells

We aimed to find novel TF perturbations capable of “rejuvenating” late passage fibroblasts. We defined rejuvenation as reversing late passage cell gene expression and phenotypes back to an earlier passage state. Our approach is depicted in the schematic diagram in Figure 1A, where each point in the high dimensional gene expression space represents one cell. Late passage and early passage wild-type (WT) cells cluster apart from each other due to their gene expression differences. Most late passage cells with TF perturbations will also cluster near WT late passage cells. However, cells with rejuvenating TF perturbations will shift towards the early passage WT cells.

**Figure 1:**
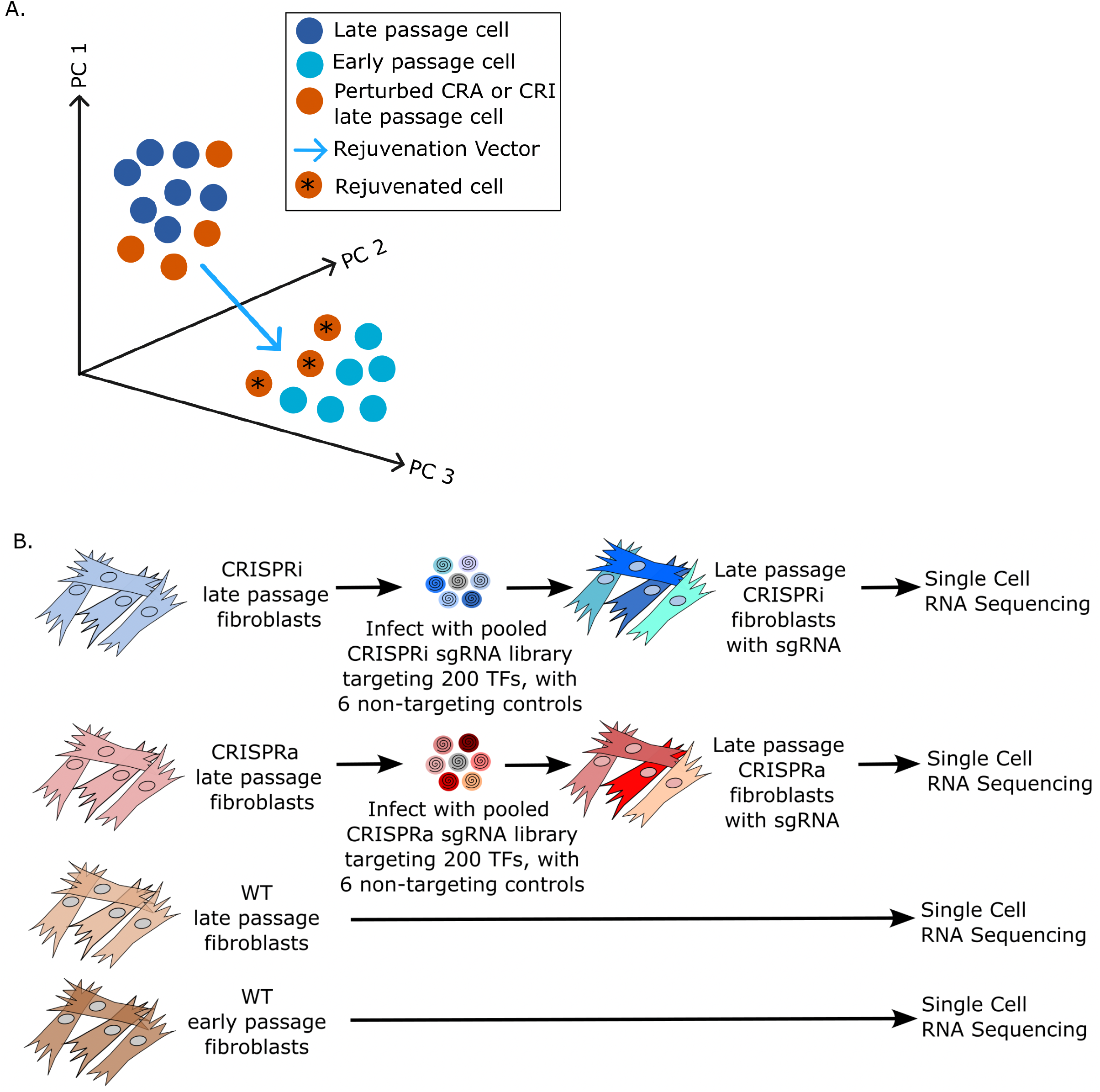
Conceptual diagram and experimental set up for discovering novel transcription factor (TF) perturbations capable of reversing gene expression in late passage skin fibroblasts back to an earlier passage state. A. Diagram of the high dimensional gene expression space indicating early passage, late passage, and perturbed late passage cells. Perturbed late passage cells that cluster close to early passage cells are considered to be “rejuvenated”. The rejuvenating TF perturbations “move” the late passage cells along the direction of the rejuvenation vector, defined as the difference between early and late passage cells. B. Experimental set up. Late passage cells expressing CRISPRa (CRA) or CRISPRi (CRI) constructs were transfected with a sgRNA library targeting different TFs for activation or repression. The gene expression state of the transfected cells was then analyzed via Direct Capture Perturb-seq to quantify the mRNA expression and identify the sgRNA in each single cell. WT early and late passage cells were assayed in parallel to define the direction of aging (or its reverse, the direction of rejuvenation).

To define the young and old states, we first measured the gene expression of passaged WT fibroblasts using single-cell RNA sequencing (scRNA-seq). Based on preliminary cell experiments and gene expression data, we categorized the cells as follows: WT cells from population doubling (PD) 1 - 19 are early passage, cells from PD 20 - 30 are middle passage, and cells from PD 30 - 39 are late passage. At approximately PD 40, cells become fully senescent. Gene expression patterns largely recapitulated previous fibroblast and cellular aging data^10,11,17^. For example, late passage cells had lower gene expression related to the cell cycle, mitochondria, proteasome, and ribosome biogenesis. On the other hand, late passage cells had more gene expression related to secretory pathways, extracellular matrix, and senescence. Using this data, we next aimed to identify candidate TFs likely playing a role in these gene expression differences.

### Identifying TF perturbations that reverse global gene expression of late passage cells back to that of earlier passage cells

There are approximately 1,500 transcription factors, cofactors, and chromatin regulators^18,19^ in the human genome. Using our lab’s TF prediction tool (see Methods), we performed a bioinformatic analysis on the scRNA-seq data to identify differentially expressed TF modules in WT passaged cells. We also performed a literature search on TFs linked to senescence and cellular aging. From these methods, we selected 200 candidate TFs to perturb (2/3 from the computational analysis and 1/3 from the literature search).

To alter the expression of the 200 candidate TFs, we created stable cell lines in late passage fibroblasts expressing either dCas9 CRISPR activation (CRA) or interference (CRI)^14,15^ to overexpress or repress TFs, respectively. Then, we infected these cells with a guide RNA (sgRNA) library targeting each of these 200 TFs ^20^, along with six non-targeting control guides, and performed Direct Capture Perturb-seq^14,21^ (Figure 1B). Direct Capture Perturb-seq is a high-throughput method for performing scRNA-seq on pooled genetic perturbation screens (using a pooled CRISPR sgRNA library). It quantifies the gene expression changes associated with a particular perturbation by simultaneously capturing the mRNAs and the sgRNA sequences in single cells. We could thus identify TF perturbations that reversed the gene expression changes caused by the replicative aging. To ensure the robustness of gene activation and repression, we targeted each TF with two distinct sgRNAs (built in the same construct) with the highest efficacy based on a previously developed library design rules^20,21^(see Methods).

For each TF perturbation, we calculated the gene expression differences between late passage cells with a TF perturbed versus late passage cells with non-targeting (NT) sgRNAs (control cells). We then compared these gene expression differences with those between WT (no CRISPR construct) late passaged cells and WT early passaged cells. TF perturbations with a significant negative correlation (as measured by the Pearson correlation coefficient r-rejuvenation) indicated the perturbation reversed the gene expression changes due to replicative aging. From the 200 overexpressed and repressed TFs, we identified more than a dozen TF perturbations with strong negative r-rejuvenation (see the top 15 hits for CRA and CRI in Tables 1 and 2, and examples of the global gene expression changes in Figure 2), suggesting that late passaged fibroblast cells may be rejuvenated by targeting these TFs.

**Table 1.** Top 15 CRISPRa (CRA) TF perturbations, ranked by r-value, including the log 2 fold change (log2fc) of the TF itself and the p value for the log2fc of the TF. Those in red boxes are the TFs we tested extensively with cell and molecular phenotyping.

| gene | r value | log2fc | p value |
| --- | --- | --- | --- |
| DLX6 | -0.57 | 0.53 | 2.00E-03 |
| E2F3 | -0.53 | -0.18 | 0.46 |
| FOXM1 | -0.47 | 2.40 | 1.70E-04 |
| FOSL1 | -0.45 | 2.07 | 5.55E-18 |
| NFATC4 | -0.41 | 0.46 | 0.12 |
| MYC | -0.40 | 1.50 | 6.00E-07 |
| STAT4 | -0.39 | 5.30 | 4.36E-14 |
| GATA3 | -0.38 | 3.30 | 5.18E-15 |
| EZH2 | -0.36 | 5.10 | 2.62E-29 |
| PLAU | -0.35 | 4.50 | 5.04E-11 |
| HSF2 | -0.35 | 0.18 | 0.48 |
| PAX4 | -0.33 | 5.50 | 5.05E-32 |
| NKX2-2 | -0.32 | 0.15 | 0.16 |
| SIM2 | -0.32 | 3.41 | 2.24E-13 |
| VAX1 | -0.30 | 2.56 | 2.78E-08 |

**Table 2.** Top 15 CRISPRi (CRI) TF perturbations, ranked by r-value, including the log 2 fold change (log2fc) of the TF itself and the p value for the log2fc of the TF. Those in red boxes are the TFs we tested extensively with cell and molecular phenotyping.

| gene | r value | log2fc | pval |
| --- | --- | --- | --- |
| EGR1 | -0.55 | -1.82 | 9.87E-10 |
| ZFX | -0.51 | -0.9 | 5.57E-10 |
| ATF4 | -0.48 | -1.85 | 1.25E-09 |
| VEZF1 | -0.47 | -1.73 | 7.07E-14 |
| MAZ | -0.47 | -1.28 | 2.53E-07 |
| SOX2 | -0.47 | 0.00 | 0.31 |
| PAX8 | -0.44 | 0.00 | 0.91 |
| ZFHX3 | -0.43 | -0.11 | 0.63 |
| ATG4C | -0.42 | -0.62 | 9.36E-09 |
| PLAU | -0.41 | -2.62 | 2.01E-23 |
| ATG5 | -0.41 | -2.3 | 3.93E-13 |
| HMGB1 | -0.41 | 0.07 | 0.53 |
| STAT3 | -0.41 | -2.51 | 8.30E-12 |
| GATA2 | -0.40 | 0.52 | 0.09 |
| KLF4 | -0.39 | -0.76 | 1.24E-05 |

**Figure 2.**
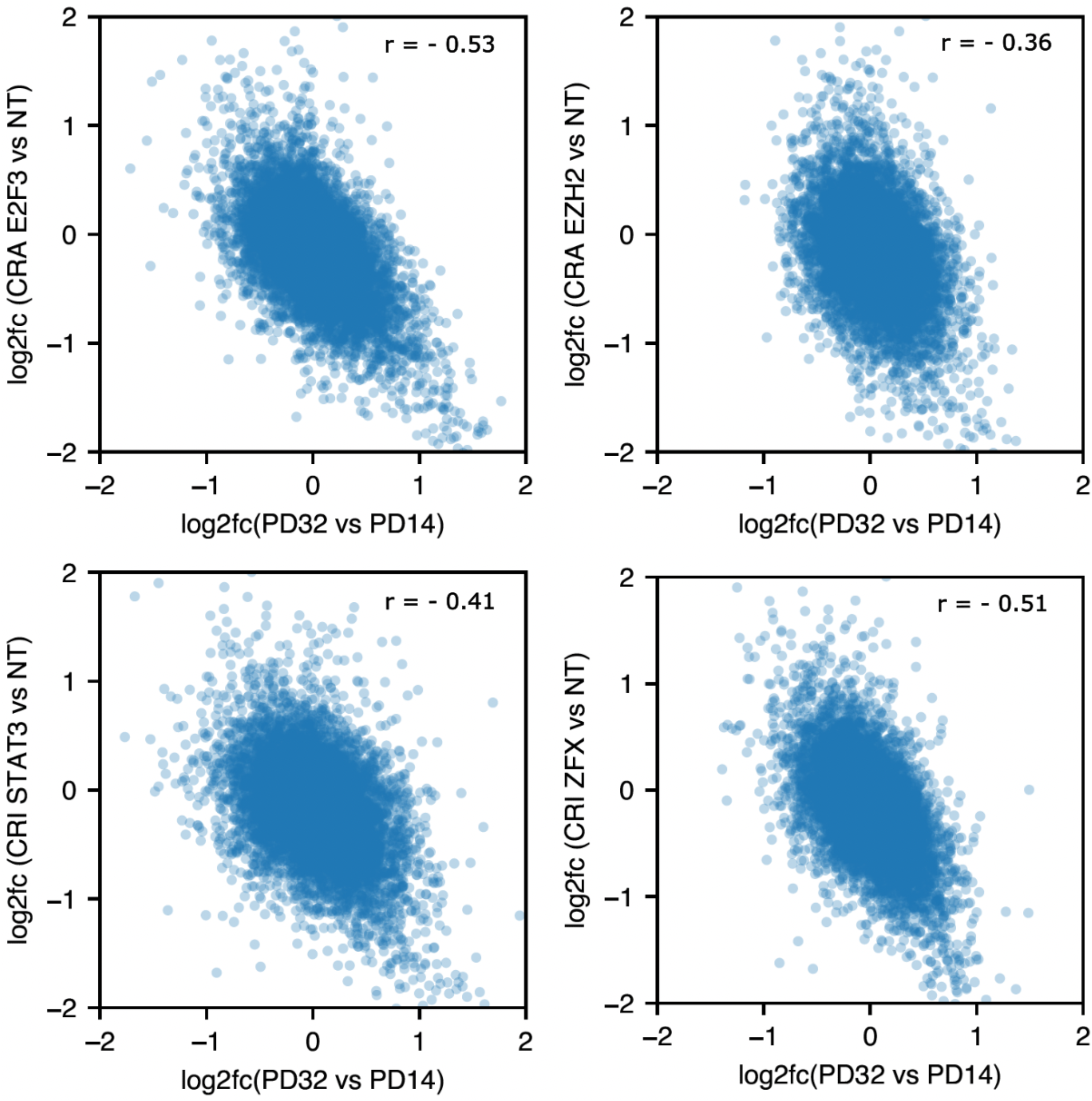
Examples of TF perturbations that reversed gene expression in late passage cells back towards an earlier passage state. Correlation plots comparing gene expression changes between late passage and early passage WT cells to that between late passage cells with a TF perturbation and those with the NT control. Shown are CRA E2F3, CRA EZH2, CRI STAT3, and CRI ZFX, the four TF perturbations that we subsequently validated with cellular and molecular phenotyping. TF perturbations with a significant negative correlation (as measured by the Pearson correlation coefficient r-value) indicated that the TF perturbation reversed gene expression changes due to replicative aging.

To gain a global perspective on how the transcriptional landscape of late passage cells changed due to these TF perturbations, we performed a TF module analysis by applying a previously developed computational method called SCENIC^22^ to our scRNA-seq data. This analysis revealed that several top TF perturbations caused similar gene expression changes in the late passage cells, even though the TFs themselves are from diverse upstream pathways (Figure 3A).

**Figure 3.**
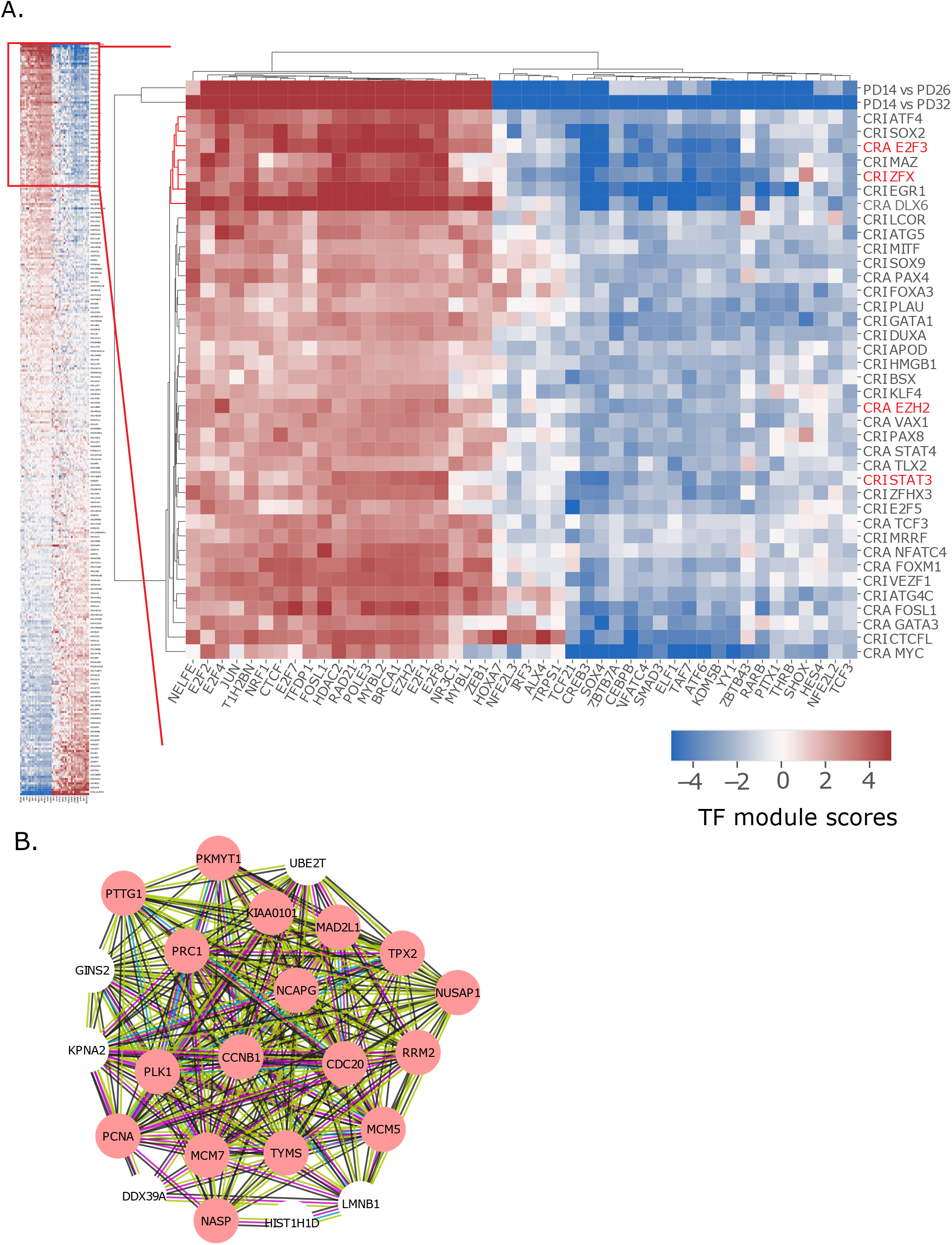
Transcription factor module analysis revealed that rejuvenating TF perturbations drive similar downstream gene expression changes. A. TF perturbations (rows) were clustered by the AUC t-test scores for selected TF modules (columns) from the SCENIC analysis. Only the modules differentially expressed between WT PD14 (early passage) and PD32 (late passage) (|t-test score| > 5) were shown. A positive (red) module score means the genes in that TF module were more expressed in the TF perturbation compared to the NT control, or earlier passage cells compared to later passage cells, and a negative (blue) module score means the genes in that TF module were less expressed in the TF perturbation or earlier passage cells. The AUC t-test score is derived from the AUC score for individual cells from the SCENIC analysis (see Methods). B. Interaction network analysis (using string-db) of commonly up-regulated genes within a cluster of TF perturbations in A (CRA E2F3, CRA DLX6, CRI ZFX, CRI EGR1, CRI MAZ, CRI SOX2, and CRIATF4). The red color indicates genes in the cell cycle gene ontology category. The lines connecting genes represent known interactions (light blue and pink), predicted interactions (green, red, dark blue) and other potential connections (yellow, black, purple).

We further investigated whether specific genes, and not just TF modules, overlapped as well. In fact, within the top 100 most significantly up-regulated genes for seven TF perturbations that clustered together in Figure 3A (CRA E2F3, CRA DLX6, CRI ZFX, CRI EGR1, CRI MAZ, CRI SOX2, and CRI ATF4), 23 genes were up-regulated in at least five of seven TF perturbations (Figure 3B)^23^. These data indicate (1) passaged fibroblasts have tightly interconnected gene regulatory networks and (2) perturbing different TFs can lead to similar gene expression outcomes, possibly via transcriptional cascades through the networks.

### Validating top TF hits from the Perturb-seq screen through cellular and molecular phenotyping of aging hallmarks

To test whether the top TF perturbations from our Perturb-seq screen indeed rejuvenated late passage cells, we performed comprehensive cellular and molecular phenotyping of various aging hallmarks^11^ in cells with a specific TF targeted. We identified four novel TF perturbations that consistently rejuvenated diverse aging hallmarks: CRA (overexpression) of EZH2 or E2F3, and CRI (repression) of STAT3 or ZFX. E2F3 is largely involved in regulating cell cycle progression from G1 to S phase^24^. EZH2 is a methyl-transferase best known for being a catalytic subunit of the polycomb repressive complex 2 (PRC2)^25^ but has other roles outside the PRC2 complex^26–28^. STAT3 is a member of the STAT family that forms part of the JAK-STAT signaling cascade and plays an important function in immune/inflammatory response^29^. ZFX is relatively poorly understood TF^30^, but it has links to stem cell renewal^31,32^. We selected these four TF perturbations initially because they had large negative r-rejuvenation scores and the strongest phenotypes in cell cycle gene expression^33^ and cell growth assays. Excitingly, targeting any one of these four TFs alleviated diverse cell aging phenotypes, confirming their rejuvenating effect beyond the transcriptional program.

Decreased cell proliferation and increased cellular senescence are two defining hallmarks of replicative aged fibroblast cells^10,11^. We found that perturbing any of the four TFs leads to significantly more cell division, as measured via immunofluorescence of KI67, a common cell division marker^34^ (Figure 4A, 4B), and by cell cycle analysis^33^ of the gene expression data (Figure 4C). For all four TF perturbations, the cell division rate returned to that of a middle passage state (about 12 - 14 PDs earlier). CRA E2F3 and CRA EZH2 caused significant decreases in cellular senescence, as quantified by senescence associated beta-galactosidase staining^35^(Figure 5A, 5B). In addition, all four TF perturbations had lower expression of senescence associated genes p21, TIMP1, and TIMP2^36^, while late passage cells expressed more (Figure 6 A-C). Thus, targeting any one of these four TFs in late passage cells caused more cell division and less senescence.

**Figure 4.**
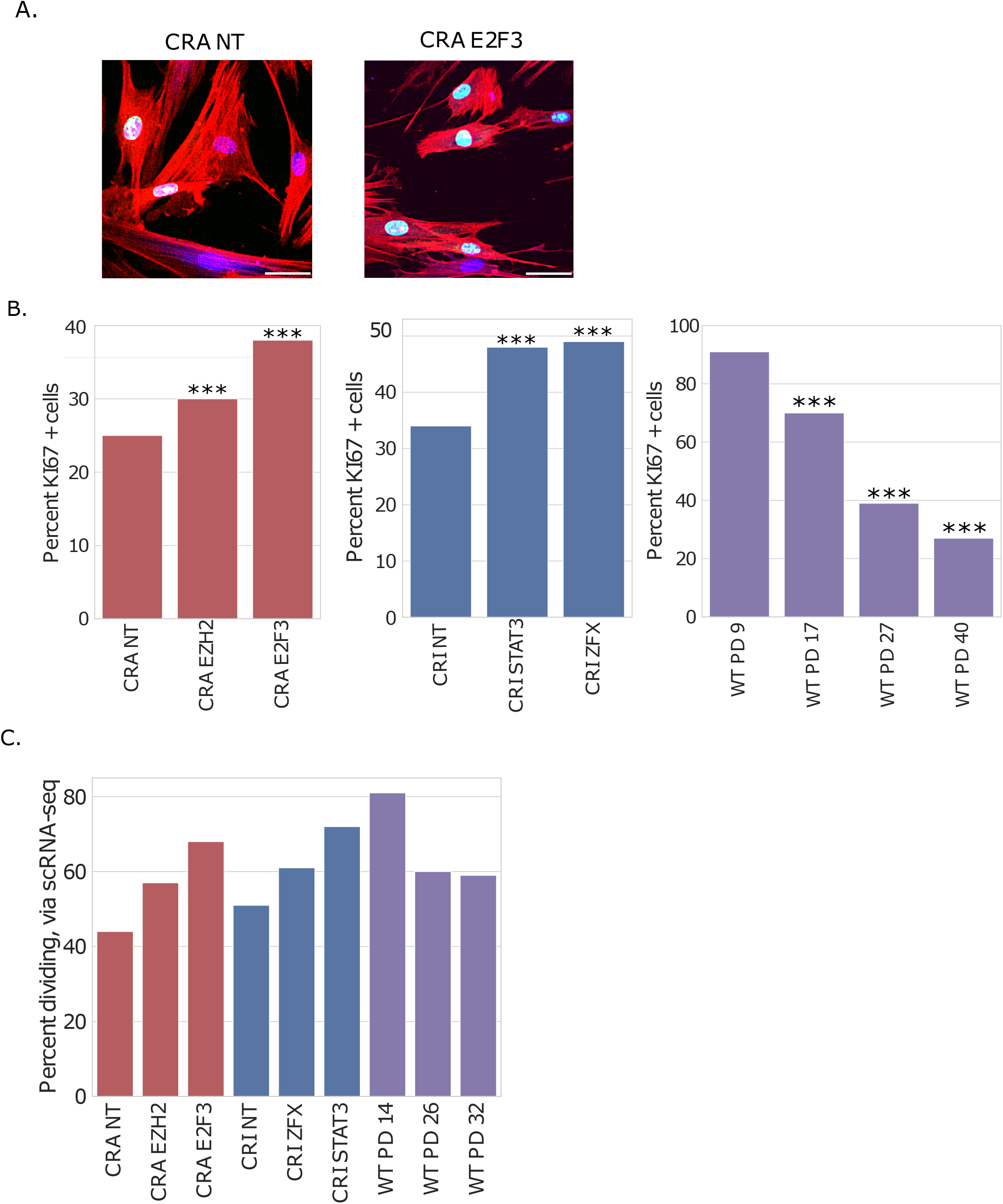
TF perturbations increased the cell division rate. A. KI67 microscopy of CRA NT and CRA E2F3 cells. Blue is Hoechst staining the nucleus, red is phalloidin staining actin, and green is KI67; 50 uM scale bar. B. Percent KI67 positive cells for CRA and CRI perturbed cells and WT passaged cells ranging from early to late population doubling (PD); N >1500 per sgRNA, and N >700 per WT PD; statistical significance is calculated by binomial distribution, relative to PD 9 for WT and NT for CRA and CRI. Data from experiments performed on different days was normalized and combined as described in Methods. Similar data normalization was performed for other figures; *p <0.05, **p <0.01, ***p <0.001. C. The percent of cells in S, G2, or M phase of the cell cycle, as measured via single cell RNA sequencing analysis.

**Figure 5.**
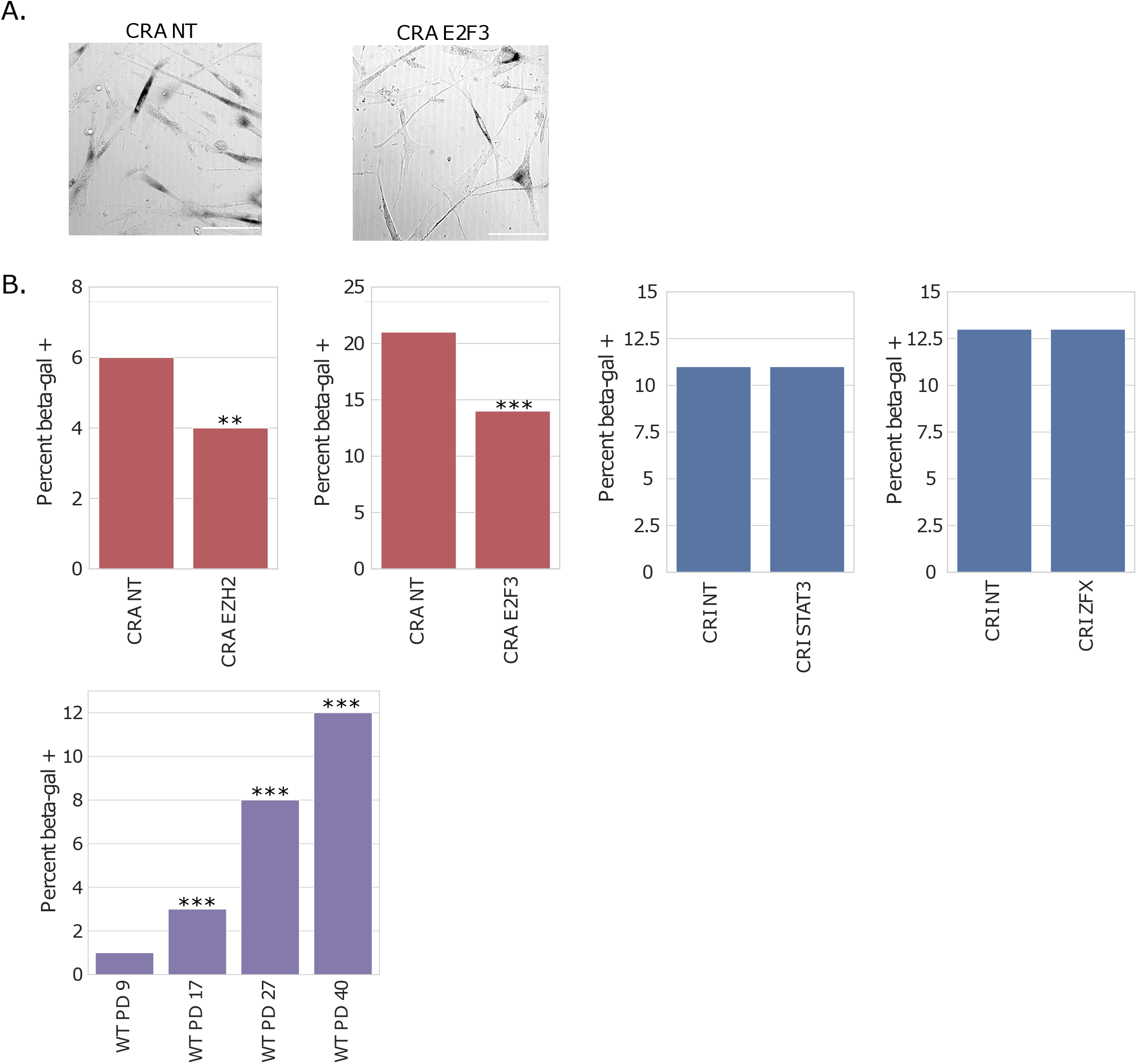
TF perturbations CRA E2F3 and CRA EZH2 decreased the number of senescent cells. A. CRA NT and CRA E2F3 cells stained for beta-galactosidase (beta-gal), imaged in brightfield, 100 uM scale bar. Dark cells are beta-gal positive. B. Percent beta-gal positive cells for CRA, CRI, and WT passaged cells; N >700 cells per sgRNA, N >400 for each WT PD. Statistical significance is calculated by binomial distribution, relative to NT for CRA and CRI and PD 9 for WT. *p <0.05, **p <0.01, ***p <0.001.

**Figure 6.**
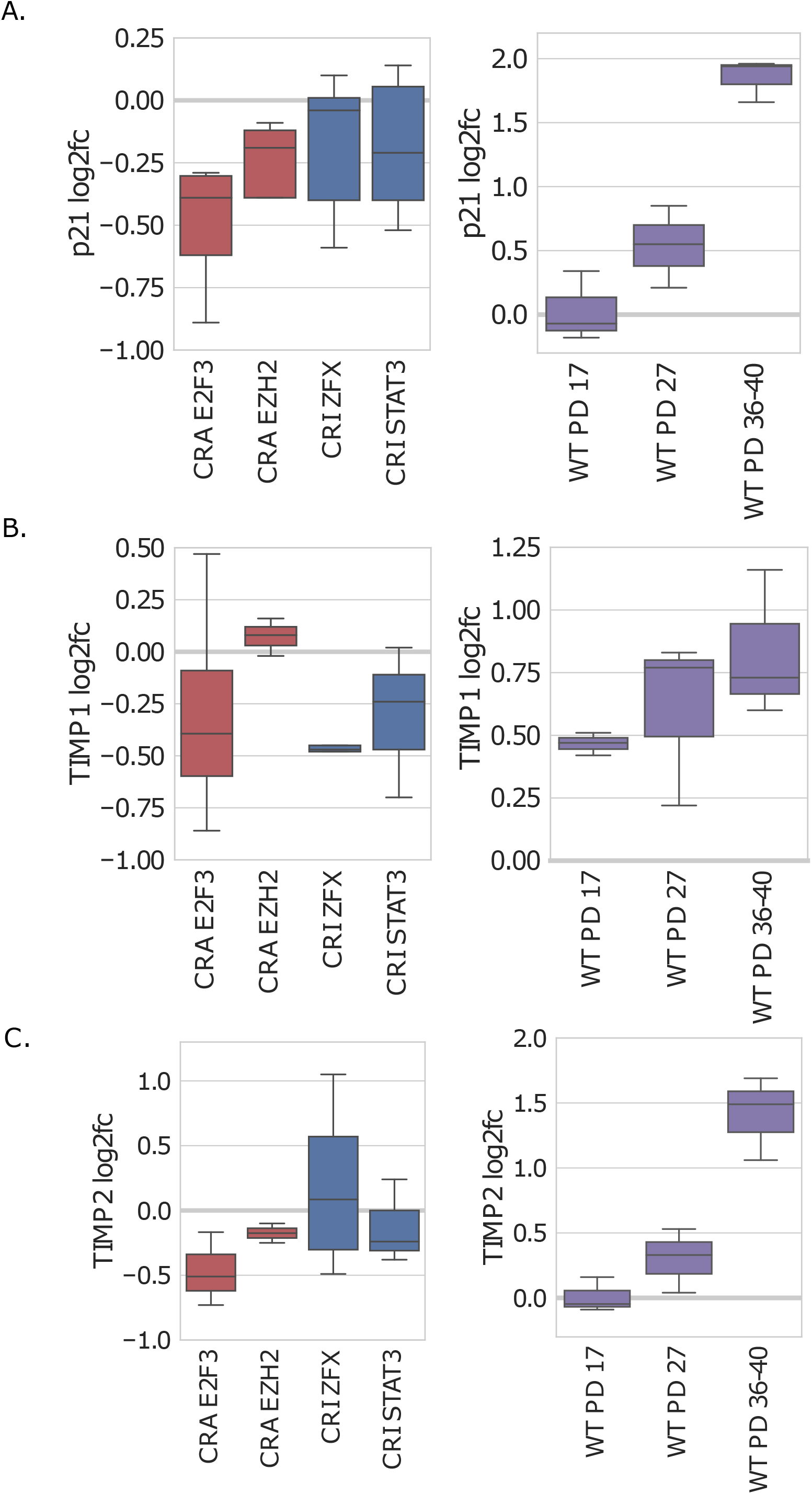
TF perturbations decreased the expression of common senescence associated genes. Using quantitative PCR (qPCR), we measured changes in the expression of A. p21 (CDKN1A), B. TIMP1, and C. TIMP2 for CRA and CRI is relative to NT controls, and the log2fc for WT is relative to early passage cells

Loss of proteostasis is a key contributor to several diseases and aging^37^. All four TF perturbations significantly increased proteasome expression, reversing the pattern seen in late passage cells (Figure 7A). In three of the four TF perturbations, there was also significantly increased proteasome activity, as measured through a fluorescence-based cleavage assay (Figure 7B).

**Figure 7.**
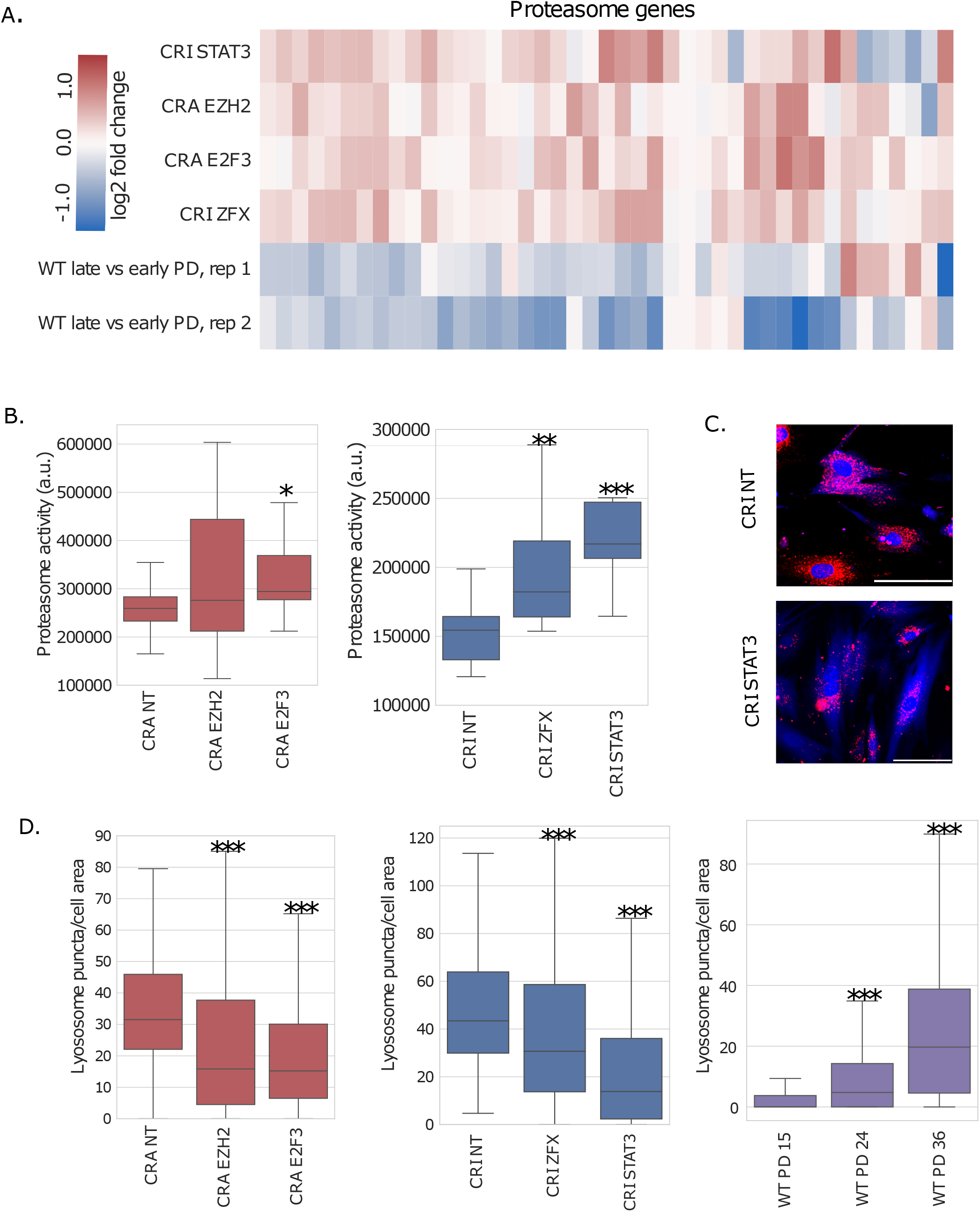
TF perturbations improved proteostasis. A. Cluster heatmap of all proteasome genes, colored by log2 fold change (log2fc). In CRA and CRI cells, the log2fc is relative to NT cells; for WT, the log2fc is relative to early passage cells (WT PD 32 versus PD 14). B. Proteasome activity for CRA and CRI TF perturbations; significance was calculated by a Wilcoxon rank-sum test, comparing TF perturbations to NT. *p <0.05, **p <0.01, ***p <0.001. C. CRI NT and CRI STAT3 cells stained with LysoTracker Red; blue is BFP from the sgRNA construct and labels the cytoplasm; 100 uM scale bar. D. Quantification of lysosome puncta per cell area, as measured with LysoTracker Red. N >180 cells per sgRNA, and N > 330 cells per WT PD. Significance was calculated by a Wilcoxon rank-sum test, comparing TF perturbations to NT and WT later PDs to WT PD 15. *p <0.05, **p <0.01, ***p <0.001.

Lysosomes also play an important role in proteostasis. Interestingly, there were significant decreases in total lysosome puncta per cell area in all four TF perturbations, while late passage WT cells had significantly more lysosome puncta per cell area than early passage cells, as measured by the Lysotracker staining (Figure 7C and 7D). An early electron microscopy study found the lysosomes of serially propagated human fibroblasts gradually transform to residual bodies (containing undigested materials), and these bodies increase in number and size^38^, reflecting degeneration of lysosomal function^39^. Thus, it is likely the Lysotracker stained those residual bodies, and the decrease of the number of puncta by the TF perturbations indicated improved lysosomal function. Overall, these four TF perturbations significantly rejuvenated proteostasis in these late passage fibroblasts.

Mitochondrial dysfunction is another conserved hallmark of aging^11^. Mitochondria become less functional and mitochondrial genes are less expressed in old cells and late passage cells^10,11^. In all four of our top TF perturbations, there were significant increases in mitochondrial and Krebs cycle genes, reversing the pattern seen in late passage WT cells (Figure 8A). We assayed mitochondria function by measuring the mitochondrial membrane potential, using the TMRE (tetramethylrhodamine, ethyl ester) membrane potential marker. CRA EZH2 had a significant increase in mitochondrial membrane potential (Figure 8B and 8C). CRA MYC was a positive control for increased mitochondrial membrane potential, given MYC’s known roles in mitobiogenesis^40^. CRA E2F3 had slightly less mitochondrial membrane potential. The other TF perturbations did not have significant changes in mitochondrial membrane potential.

**Figure 8.**
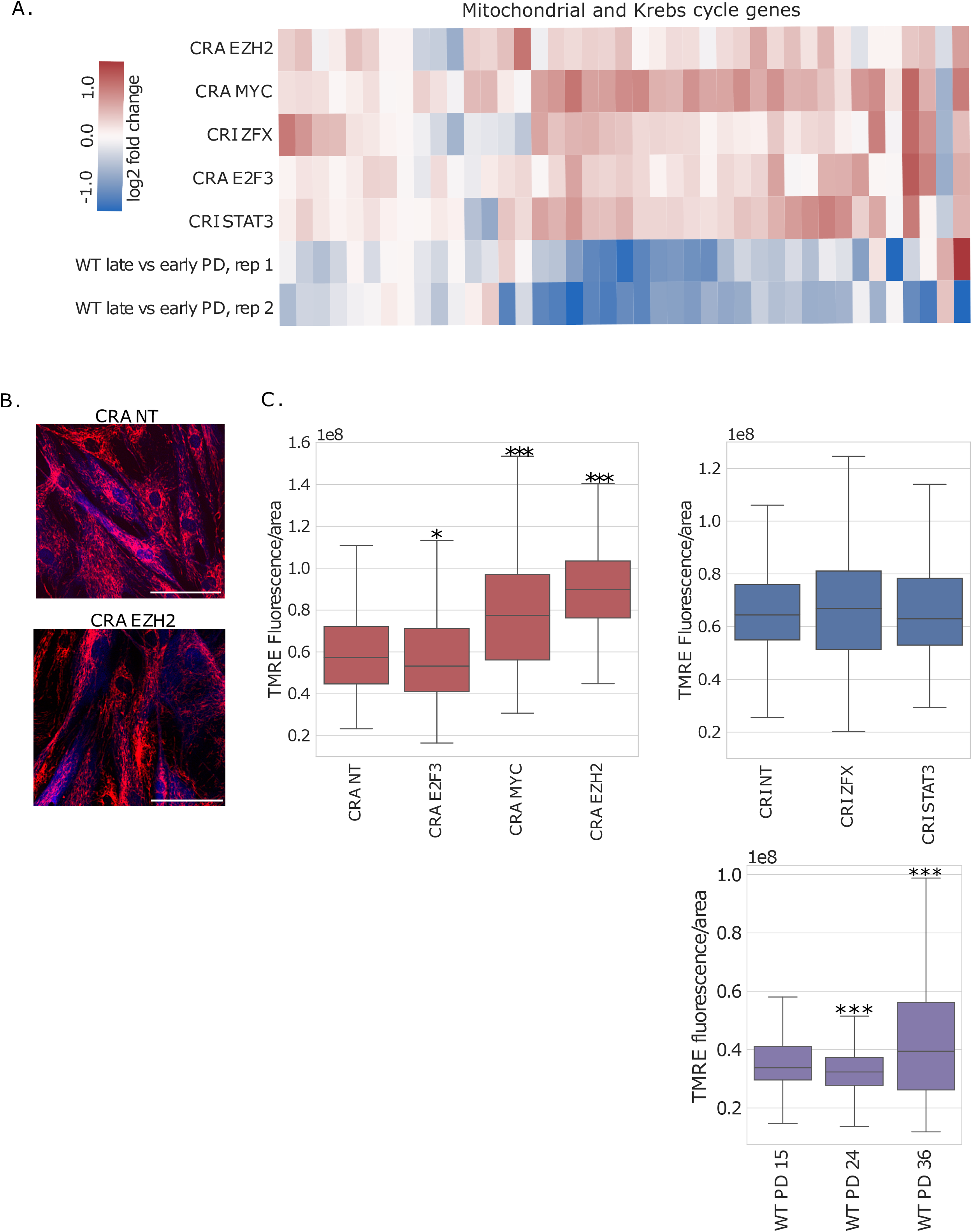
TF perturbations enhanced mitochondrial function. A. Cluster heatmap of mitochondrial genes (all “MT” mitochondrial genes except those encoding tRNA) and Krebs cycle genes, colored by log2 fold change (log2fc). In CRA and CRI cells, the log2fc is relative to NT cells; for WT, the log2fc is relative to early passage cells (WT PD 32 versus PD 14). B TMRE (tetramethylrhodamine, ethyl ester) mitochondrial membrane potential stain of CRA NT and CRA EZH2 cells; brighter red indicates higher membrane potential; blue is BFP from the sgRNA construct and labels the cytoplasm; scale bar is 100 uM. C. Quantification of the TMRE membrane potential stain. Significance was calculated by a Wilcoxon rank-sum test, comparing TF perturbations to NT and WT later passages to WT PD 15. N >300 cells per sgRNA, N >465 cells per WT PD. *p <0.05, **p <0.01, ***p <0.001.

Telomeres, protective caps at the ends of chromosomes, get progressively shorter every time a cell divides^41^, and late passage cells have shorter telomeres than early passage cells^42^. Overexpressing telomerase (TERT) increases telomere length, allowing cells to divide regardless of their usual Hayflick limit^43,44^. Thus it is essential to ask whether the rejuvenating effect of the four TFs depends on the activation of telomerase and increased telomere length. We compared gene expression of our top TF perturbations to previously published data on dermal skin fibroblasts overexpressing TERT^45^. None of the TF perturbations have similar gene expression changes to TERT overexpressing fibroblasts (Figure 9A). Furthermore, TERT mRNA itself was never expressed enough to be measured in our scRNA-seq experiments (data not shown). We next measured the relative length of telomeres in late passage cells with the TF perturbations, NT controls, and WT passaged cells through qPCR analysis^46^. While we did see a progressive decrease in telomere length in passaged cells, there was no change among TF perturbations (Figure 9B). Therefore, the TF perturbations did not affect telomere length, and their rejuvenation phenotypes are independent of the telomerase and telomere axis.

**Figure 9.**
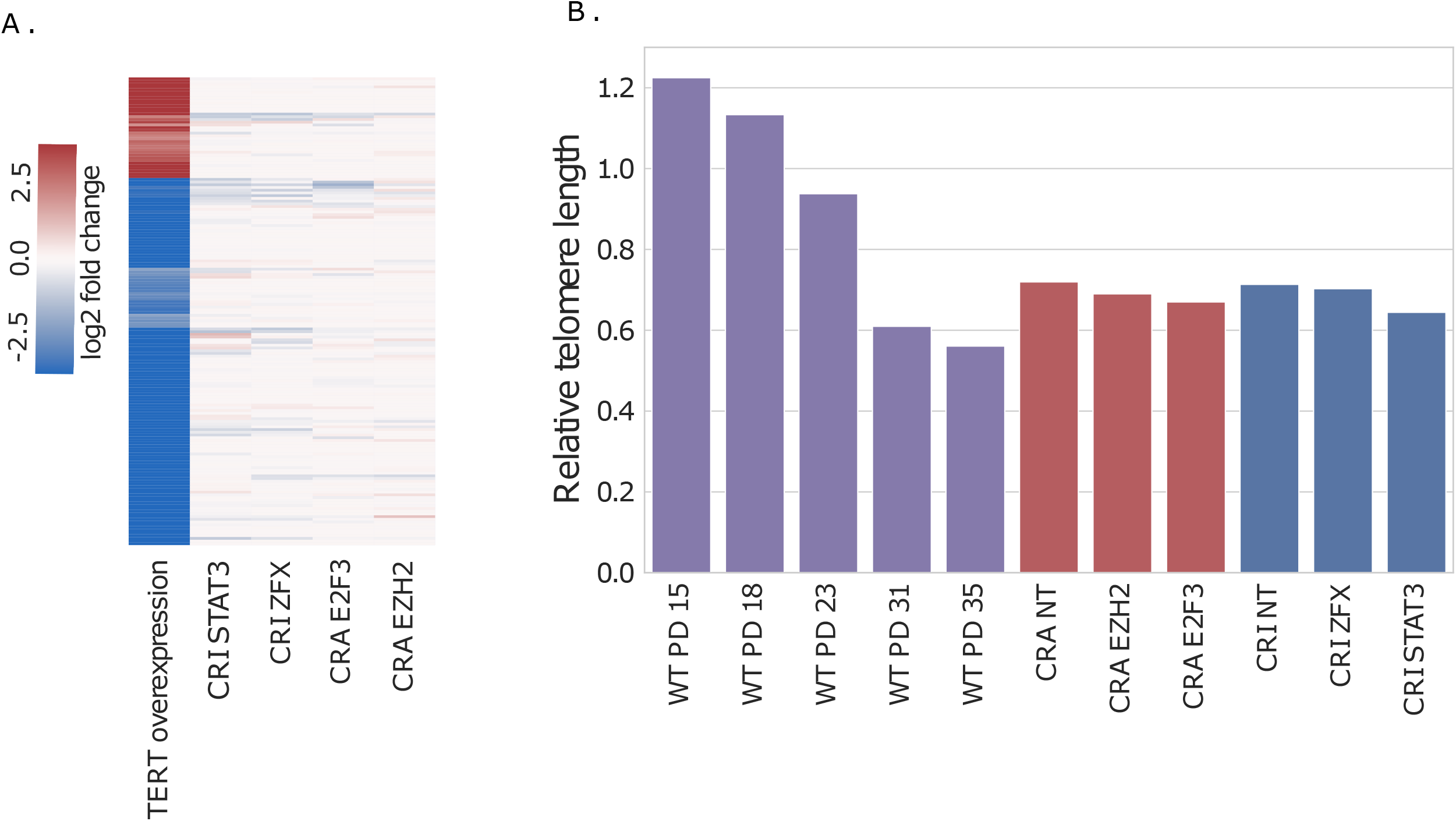
TF perturbations and their effects on late passage cells are independent of telomeres. A. Differentially expressed genes when TERT was overexpressed in human skin fibroblasts (previously published data), colored by log2 fold change (log2fc). In CRA and CRI cells, the log2fc is relative to NT cells; for TERT overexpression, the log2fc is relative to untransformed primary fibroblasts. B. Relative telomere length, as determined through qPCR analysis, for WT passaged cells and CRA and CRI TF perturbations.

As rejuvenation of replicatively aged cells necessarily increases their proliferation, an important question to ask is whether the TF perturbations make the late passage cells behave like cancer cells. In Yamanaka factor rejuvenation experiments, cells become cancerous if the four TFs are turned on too much or for too long^3,47^. We compared the gene expression of our top TF perturbations to that of previously published data on dermal skin fibroblasts transformed into cancer cells across several steps^45^. In that work, cells were first immortalized with TERT overexpression, then transformed with SV40 large-T antigen, and finally metastasized with oncogenic H-Ras (RASG12V)^45^. None of our TF perturbations had similar gene expression changes to transformed or metastasized skin fibroblasts (Figure 10A and 10B). When looking at genes commonly differentially expressed across seven types of cancer^48^, there were some genes more expressed in our top TF perturbations (Figure 10C). But, all those genes are cell cycle related^45^. Because our top TF perturbations also caused more cell division, this overlap was not surprising. In all our experiments, the TF perturbations maintain the normal growth rate of primary fibroblast cells. Even when we extended our experiment and perturbed these TFs for over twice as long as our typical experiments, cells still grew at middle passage cell rates (data not shown).

**Figure 10.**
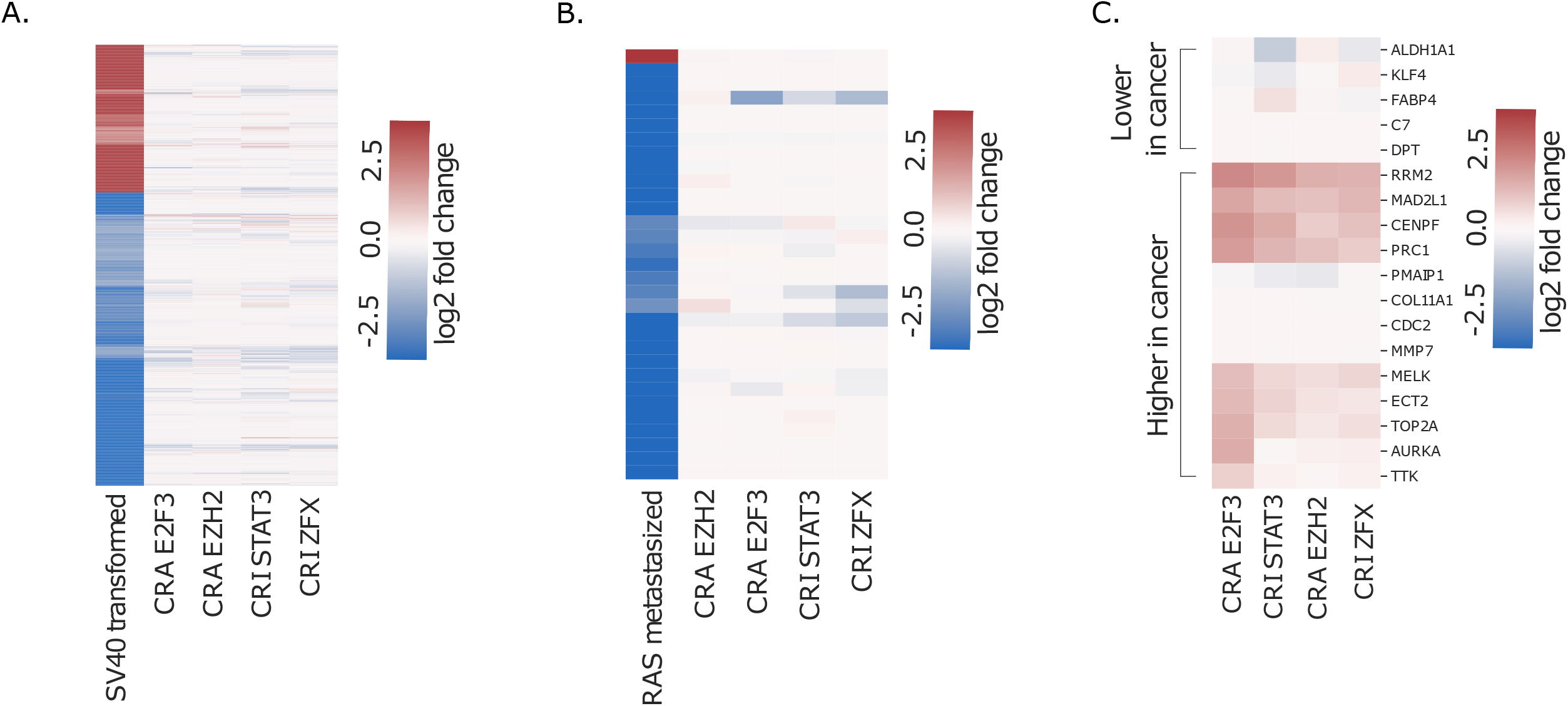
TF perturbations did not lead to cancer-like gene expression. A. Differentially expressed genes when SV40 large-t antigen was overexpressed in skin fibroblasts (previously published data). In CRA and CRI cells, the log two fold change (log2fc) is relative to NT cells; for SV40 cells, the log2fc is relative to TERT overexpressing fibroblasts. B. Differentially expressed genes when oncogenic H-Ras (RASG12V) was introduced to skin cells (previously published data). In CRA and CRI cells, the log2fc is relative to NT cells; for cells with Ras overexpression, the log2fc is relative to SV40 cells. C. Commonly differentially expressed genes from seven cancer types. In CRA and CRI cells, the log2fc is relative to NT cells.

Overexpressing all four Yamanaka factors at once is sufficient to cause cancer. While the four top TFs we identified do have links to cancer, literature supports that changing their expression individually does not seem sufficient to cause cancer^49–56^. STAT3 and ZFX are often overexpressed in cancer^49,52^, but in our studies, we found repressing these genes was rejuvenating in late passage fibroblasts. While our gene expression analysis suggests these TF perturbations are unlikely to cause cancer in an *in vitro* system, further experiments are necessary to fully determine their carcinogenic potential.

We also analyzed other aging hallmarks related to genome instability (DNA damage markers) and epigenetic modifications (histone and DNA methylation), and did not find consistent changes due to the TF perturbations.

DNA damage increases as cells age, but also as cells grown in culture divide more rapidly^11,57,58^. A common way to measure DNA damage is throughγH2AX, a histone phosphorylation marker adjacent to double stranded DNA breaks^59 60^. In rejuvenation studies with the Yamanaka factors, γH2AX decreased slightly^3^. In our top TF perturbations, we saw either no change or a slight increase inγH2AX per cell (Figure S1A and B). In passaged WT cells, late passage cells did have significantly moreγH2AX puncta.

Epigenetic patterns, like histone and DNA methylation, regulate gene expression and have links to aging and senescence^11^. The DNA methylation clock correlates methylation at certain CpG islands to the actual age of organisms or PD of cells^61,62^. While the later passage WT cells did have progressively “older” methylation clock scores, none of our four TF perturbations changed the cells’ skin DNA methylation clock age ^62^(Figure S2). Thus, the rejuvenation phenotypes were not dependent on turning back the DNA methylation clock.

The global levels of histone 3 lysine 9 trimethylation (H3K9me3) and histone 3 lysine 27 trimethylation (H3K27me3) have been used as cell aging markers, but both have mixed results in various studies and organisms^63,64,65^. For example, overexpressing the Yamanaka factors in fibroblasts led to more global H3K9me3^3^; but, in a progeria model, more H3K9me3 was linked to senescence^66^. In our WT cells, late passage cells had more global H3K9me3. In the TF perturbations, global H3K9me3 levels were significantly higher in CRI STAT3 and lower in CRI ZFX. In CRA E2F3, the distribution of H3K9 global fluorescence was significantly different than in NT, although their medians were similar (Figure S3A-B). H3K27me3 levels of WT passaged cells increased significantly in later passage stages. In the TF perturbations, H3K27me3 levels were significantly higher in CRA E2F3 and significantly lower in CRA EZH2 and CRI STAT3. In the future, perhaps probing specific histone methylation sites, instead of global levels, may be more informative.

## Discussion

Using a human cell culture model of replicative aging (the Hayflick model^9,13^), we developed a high throughput screen to identify the potentially rejuvenating TFs—TFs that, when overexpressed or repressed, reprogram the global gene expression state from the late passage state back to an earlier passage state. Our approach combined a novel bioinformatic analysis to narrow down a candidate list of TFs and chromatin modifiers, followed by Perturb-seq^14,21^ to identify the potential rejuvenating factors. The top TF perturbations were then followed by extensive cell and molecular phenotyping to test the rejuvenating effects. This approach led to the successful identification of four TFs that, when overexpressed (E2F3, EZH2) or repressed (STAT3, ZFX), reversed various cell aging and senescence phenotypes. Although our experiments were only done in *in vitro* aged cells, our findings serve as a proof of concept that (1) genes outside the Yamanaka factors can reverse aging hallmarks in human cells and (2) these hallmarks can be reversed without reprogramming cells back towards a stem cell state.

Interestingly, one of our top TF perturbation hits from the screen was CRA FOXM1. A recent study showed cyclic overexpression of an N-terminal truncated form of FOXM1 delays natural and progeria aging phenotypes and extends lifespan in mice^7^. In our system, CRA FOXM1 ranked highly by the rejuvenation r-value and induced a downstream transcriptional profile similar to our newly discovered TFs (Table 1 and Figure 2). In our follow-up experiments, CRA FOXM1 did phenocopy some features seen in our top TF perturbations—fewer senescent cells, lower senescence genes expression, fewer lysosome puncta—but CRA FOXM1 did not rejuvenate cells to the same extent as our other top TF perturbations (Figure S4A-F). We suspect that these results occurred because we solely targeted the expression of endogenous FOXM1 instead of the truncated form, which is constitutively active. Because FOXM1 can rejuvenate mice, we are excited by the possibility of other TF perturbations we identified being able to rejuvenate tissues and organs *in vivo*.

Besides the four TFs we discovered, there are likely other TF perturbations which could reverse cell aging phenotypes in passaged fibroblasts. For example, CRI of ATF4 and EGR1 and CRA of DLX6 all ranked in the top three with large negative rejuvenation r-values, and their downstream transcriptional profiles clustered together with E2F3 and ZFX. Interestingly, ATF4 is a master transcriptional regulator of integrated stress response (ISR)^67^, and previous work showed that inhibition of ISR increased the survival of nematodes and improved the cognitive function of aging mice^68–71^. EGR1 plays an important role in regulating the response to growth factors, DNA damage, and ischemia^72,73^. DLX6 encodes a member of a homeobox TF gene family similar to the Drosophila distal-less gene and is much less studied. We did not pursue these factors further in this study as we decided to focus our effort on the four TFs that yielded stronger phenotypes in our initial tests based on cell cycle gene expression analyses and KI67 microscopy. However, EGR1, ATF4, DLX6 and other top hits warrant further systematic evaluation for their rejuvenating potential.

We observed that seemingly unrelated TFs led to converging downstream gene expression and cell rejuvenation signatures in late passage fibroblast cells. This observation suggests the transcriptional networks in these cells are densely connected, where targeting one node leads to similar transcriptional cascade altering the other connected nodes. It will be important to better understand the structure of the networks, and our systematic Perturb-seq data will serve as a good resource for furthering such investigation. It will also be interesting to investigate whether such convergence is seen in different models of aging and rejuvenation (e.g., different cell models or rejuvenation related treatments), and if so, whether the converged signature will overlap with what we observed in the passaged fibroblasts.

Replicative senescence in human fibroblast cells is linked to telomere attrition, and overexpressing telomerase has been shown to extend the proliferative capacity of the cells beyond the Hayflick limit^43,44,74^. The four TF perturbations we identified rejuvenated late passage cells without increasing telomere length or obviously increasing telomerase expression. And, the transcriptional changes induced by the TF perturbations did not resemble that of telomerase overexpression, suggesting that the TF perturbations caused distinct changes from telomerase overexpression. Our data support previous findings that, as the cells are continuously passaged, they progressively have more aging phenotypes and shorter telomeres. However, the aging phenotypes seem to be decoupled from telomeres until their length becomes critically short, at which point massive genome instability and cell senescence happen throughout the cell population.

The finding of multiple solutions to cellular rejuvenation will likely increase the probability of developing safe rejuvenation therapies. Currently, rejuvenation therapeutics companies are largely focused on the Yamanaka factors, where the dose and schedule of induction must be carefully controlled to avoid dedifferentiation or cancer. We observed that these four TF perturbations did not change the cell identity. In addition, the cells’ transcriptional profiles did not resemble that of cancerous cell transformations. These data point towards the possibility of rejuvenation while maintaining cell identity.

There is significant therapeutic potential in cell rejuvenation, but how to do so effectively and safely remains a challenge. To move from the lab bench towards a therapeutic, finding small molecules which cause similar gene expression changes as rejuvenating TF perturbations would be beneficial. Our finding of individual rejuvenating TF perturbations (instead of TF combinations) sets the stage for testing small molecule compounds for rejuvenation, where fluorescent transcriptional reporters of the TFs could be constructed and used for high throughput screens of small molecule libraries.

We believe the systematic approach we developed in this study can be applied to a general class of problems: searching for TFs which transform the cellular state into a predefined state with desired properties. In the context of aging and rejuvenation, the desired goal is to transform late passage cells back to early passage cells. In the context of disease, the desired goal could be to transform “diseased” state to a “healthy” state, e.g., in a cell culture model of Alzheimer’s disease. Similarly we can start by characterizing the difference between the two states (such as “diseased” and “healthy”), using bioinformatic analysis to identify a list of candidate TFs, and performing Perturb-seq to identify the relevant TF perturbations.

A major limitation of our findings in this study is that the TFs were identified from one specific cell culture model of aging: the passaged human skin fibroblast. Further work is needed to test whether the identified TFs are able to rejuvenate fibroblast cells aged *in vivo*, other aged post mitotic cells, or large-scale systems like tissues, organs, or organisms.

## Contributions

C.D. and H.L. conceived and designed the experiments. J.S. and C.D. conducted the experiments. J.Z. performed the bioinformatics analyses. M.M. performed the differentially expressed transcription factor module analysis using the tools developed in the Li lab. J.L. performed the telomere length measurements. J.S. and J.Z. prepared the figures. J.S. and H.L. wrote the manuscript. All the authors proofread the manuscript.

## Acknowledgements

We thank Drs. Eric Chow, Barbara Panning, Luke Gilbert, Aimee Kao, and Peter Walter for their advice and feedback. We thank Dr. Elizabeth Blackburn for her lab’s help performing in the telomere length assay. We thank the members of the Jonathan Weissman lab for their advice and reagents. This work was supported by NIH grants 1R21AG064357, 1R21AG071899, and 1R01AG058742, and by a CZ Biohub Investigator award and a BARI investigator award to H.L.

## Materials and Methods

### Plasmids and sgRNA

All the CRISPR related plasmids were gifts from the Jonathan Weissman lab (pMH0001, pJKNp44, pJR89, pJR85, pMJ114, pMJ117, and pMJ179). sgRNA were assembled as previously described^14^. In follow-up cell aging hallmark assays, pMJ117 was used, although any of the three pMJ sgRNA backbones would have been equally valid to use.

For dual sgRNA production, previous protocols were followed, with the slight modification of changing one digestion enzyme (see citation’s supplementary note 4)^21^. Briefly, dual-guide libraries were created by PCR amplifying pooled oligonucleotides. These oligonucleotides and pJR85 were digested with BstXI/BlpI and ligated together. Then, this new intermediate plasmid and pJR89 were digested with Esp3I. The resulting piece from pJR89 was ligated into the intermediate pJR85. The final plasmids were validated via sequencing.

### Lentivirus production

Lenti-X 293T (Lx293T) were used for lentivirus production. Lx293T cells were grown in Dulbecco’s modified eagle medium (DMEM) supplemented with 10 % FBS and penicillin-streptomycin. Lentivirus was made by transfecting Lx293T with standard packaging vectors and TransIT-LT1 Transfection Reagent (Mirus, MIR 2306). Viral supernatant was harvested two days after transfection, filtered through a 0.45 um filter, and either added directly to target cells or frozen in aliquots at -80 °C.

### Cell culture and CRISPRa and CRISPRi cell lines

Neonatal primary skin fibroblasts were purchased from ATCC (PCS-201-010) and cultured in ATCC’s Fibroblast growth kit with low serum (PCS-201-041), with phenol red (ATCC, PCS-999-001) and penicillin-streptomycin (ATCC, PCS-999-002). These fibroblasts were passaged for almost one year, during which they were split about 1:2 or 1:4 at about 80 - 90 % confluency. Population doublings were determined using a standard method, by which we compared the number of cells plated initially to the number of cells at the next passage. Cells were frozen in normal medium plus 10 % DMSO for long-term storage in liquid nitrogen.

Stable cell lines in passaged fibroblasts expressing either CRISPRi (CRI; pMH0001) or CRISPRA (CRA; pJKNp44) were created by infecting either the CRA or CRI lentiviral particles into early passage fibroblasts. These vectors have a BFP tag, and thus we sorted cells for purity by BFP on the BD FACSAria2. CRA and CRI cell lines were passaged until they were late passage; BFP fluorescence and CRA/CRI activity was maintained across all population doublings. Next, sgRNA lentiviral particles were infected into CRA and CRI cells at the desired population doubling at an MOI of ∼0.3. For all experiments using sgRNA, CRA or CRI cells were infected with the sgRNA lentiviral particles, recovered for two days, selected for purity using puromycin for 2-3 days (2μg/mL), recovered for an additional 2 days, and then used for experiments. The sgRNA vector had a BFP tag as well, and this one was significantly brighter than the CRA/CRI vectors’ BFP. The puromycin selection led to 90 - 100% purity, which we could visualize by the very bright BFP signal from the infected cells.

### Real-time quantitative polymerase chain reaction (qPCR)

Total RNA was isolated using the RNeasy Plus Mini Kit (Qiagen, 74134). RNA was converted to cDNA using the SuperScript IV (Invitrogen, 18090050) standard protocol. Twenty uL qPCR reactions were prepared with 10 uL of the KAPA SYBR FAST Universal MasterMix (Roche, KK4602), 5 uL cDNA (representing 5 - 20 ng RNA per reaction), and 5 uL of forward and reverse primers mixed together 1:1 at 0.8 uM each. Three technical replicates were run for every sample. These reactions were run on the LightCycler 480. Relative expression of each gene (ΔCt) was measured using beta-actin as a control gene. Log 2 fold change (log2fc) was calculated by finding the difference between two conditions’ conditions’ΔCts (ΔΔCts).

### Single-cell RNA sequencing and Perturb-seq

For our first round of WT passaged cell scRNA-seq, we used manufacturer’s protocols for the Chromium Single Cell 3’ Library v2 (10x Genomics, 120237) with one change; different WT passage stages were added in a pool and identified with cell membrane barcodes (MULTI-seq^75^). For the second round of WT passage cell scRNA-seq, we used the same set up, except we used the Chromium Next GEM Single Cell 5’ Library v1.1 (10x Genomics, 1000167).

For the CRA/CRI Perturb-seq experiment with dual sgRNA, the top two guides for every TF and non-targeting guides were selected from a previously derived list^20^. Dual direct capture seq was performed as described previously^21^. Briefly, CRA and CRI cell lines were infected with two pooled lentiviral libraries (one library for CRA, one for CRI) of 200 dual sgRNA vectors and three dual non-targeting vectors (6 non-targeting guides total). These cells were selected for purity with puromycin (2μg/mL), recovered for two days, and processed according to the protocol for Chromium Next GEM Single Cell 5’ Library v1.1 with slight changes^21^. All scRNA-seq libraries for WT and Perturb-seq were sequenced on a NovaSeq 6000.

### Immunofluorescence

For all microscopy experiments, on day one, cells were plated in 8 well cell culture treated microscopy slides (ibidi, 80841) so the cells would be at about 70 % confluency the next day. For immunofluorescence, on day two, cells were first fixed (4% paraformaldehyde in PBS) for 10 minutes, washed with PBS, and then blocked/permeabilized (2% Bovine Serum Albumin/0.1% Triton X in PBS) for one hour at room temperature. The wells were washed with PBS, and then primary antibodies were added (buffer: 0.5% Bovine Serum Albumin/0.1% Triton X in PBS) for one hour at room temperature. The wells were washed with PBS. Then, secondary antibodies, Hoechst 33342 (Thermo Scientific, 62249), and Alexa Fluor™ 546 Phalloidin (Invitrogen, A22283) were added to the wells and incubated for one hour at room temperature in the dark.

Finally, the wells were washed with PBS and imaged in PBS. For live-cell imaging of lysosomes and mitochondrial membrane potential, cells were plated as described above. On day two, manufacturer protocols were followed for both TMRE-Mitochondrial membrane potential staining (Abcam, ab113852) and lysosome LysoTracker™ Red DND-99 staining (Invitrogen, L7528). All microscopy quantification was done using ImageJ.

### Beta-galactosidase staining

Manufacturer protocols were followed for the Senescenceβ-Galactosidase Staining (Cell Signaling Technology, 9860). To avoid evaporation of theβ-Galactosidase stain overnight, which would lead to salt crystals precipitating out of solution, the slides were placed in a plastic container with water-soaked paper towels to create a humidity chamber. Quantification was done using ImageJ.

### Proteasome activity

Manufacturer protocols were followed for the proteasome activity assay (Proteasome-Glo Chymotrypsin-like cell-based assay, Promega, G8660). The fluorescence was measured on the Promega GloMax plate reader.

### Relative telomere length

Genomic DNA was extracted from approximately one million cells per condition (QIAamp DNA Blood Mini Kit, 51104). Relative telomere length was measured by quantitative polymerase chain reaction (qPCR), expressed as the ratio of telomere to single-copy gene abundance (T/S ratio)^76,77^. Detailed protocol can be found on the Telomere Research Network’s website (https://trn.tulane.edu/wp-content/uploads/sites/445/2021/07/Lin-qPCR-protocol-01072020.pdf). The inter assay coefficient of variation (CV) for this study is 2.7% ± 1.7%. The intraclass correlation (ICC) of duplicate DNA extraction from similar samples is 0.955 (CI: 0.914-0.977).

### Methylation clock

Genomic DNA was extracted from approximately one million cells per condition (QIAamp DNA Blood Mini Kit, 51104). The genomic DNA was then brought to the Stanford Genomics Facility, where bisulfite conversion and methylation chip experiments using the Infinium MethylationEPIC Kit were conducted. For quantification of the methylation data, the methylclock package^78^ was used, specifically the skinHorvath clock. There were technical replicates for the control samples (CRA NT, CRI NT). Due to a small fraction of CpG islands having inconsistent methylation rates between repeats, these repeats had about 15 - 30 % variability in methylation clock results. To correct for this technical variation, for each CpG island, the difference of the repeats was divided by the mean of the repeats, and only those CpGs with less than 15 % variability were kept. The more variable CpGs were filtered out, meaning they did not contribute to the methylation clock calculations. The specific CpGs filtered out for CRA NT were also filtered out for CRA EZH2 and CRA E2F3; the CpGs filtered out for CRI NT were also filtered out for CRI STAT3 and CRI ZFX. Then, the mean value for the technical repeats was calculated.

### Single-cell RNA Sequencing (scRNA-seq) Analysis

10x Genomics Cell Ranger and Scanpy33 were computational packages used to analyze scRNA-seq data. The potential rejuvenation effect of the TF perturbations was measured by how well the gene expression profile in perturbed cells mimicked the gene expression profile in the early passage cells, compared to the late passage cells. We first computed the gene expression fold changes (log2) in the late passage cells compared to the early passage cells: 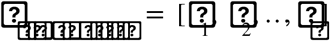 where 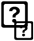 is the log2 fold-change of gene g between late passage cells and early passage cells. Then, for each TF perturbation (CRA or CRI) we computed the gene expression fold changes (log2) by comparing the cells with the guides targeting the TF and the cells with the non-targeting guides (NT): 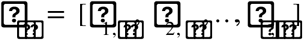 where 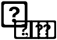 is the log2 fold-change of gene g in the perturbed cells (targeting the tf) vs the NT cells. We then computed the Pearson correlation of the gene expression vectors from each TF perturbation against 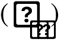 the gene expression vector of WT late passage 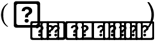. The TF perturbations with the strongest negative Pearson correlation had the most significant change in gene expression towards being like earlier passage cells.

### Differentially expressed TF module analysis for selecting initial TF candidates for the screen

Briefly, the promoter region around the transcription start site of every gene was scanned with known TF motifs with their positional weight matrices. A motif score was calculated by scanning 5000 base pairs upstream from the transcription start site for each gene^79^ and transformed into a Z score using the mean and standard deviation of the motif score for all the genes. To create a TF module, we selected all genes for a TF with a Z-score of at least 2.5; for TFs with fewer than 50 genes passing this cutoff, the top 50 genes were selected. To identify TFs related to gene expression differences between early and late passage cells, we performed a Welch’s t-test on the log2 fold change between early and late passage cells for the genes in each TF module against all other genes.

### Transcription Factor Module Analysis with SCENIC

To find the downstream transcriptional signatures of the TF perturbations, we performed TF module analysis using the SCENIC pipeline (SCENIC)^16,22^. Briefly, TF targets were inferred from the scRNA-seq data based on the co-variation between a given TF and a gene and the occurrence of the TF binding site motif in the promoter of the gene. A module activity score (AUCell score) was then computed for each module in each single cell. We then compared the AUCell scores from the perturbed cells (CRA or CRI) with the corresponding NT cells with a ranksum test to derive an AUCell t-score for the differential gene expression between the TF-perturbed and the NT cells. Similar calculations were done for WT young versus old cells.

### Statistical analysis for cellular assays

Experiments were conducted in at least three biological replicates for all cell assays for TF perturbations and NT control. Sometimes biological replicates (sub-experiments) were done on separate days. Because of slight differences in staining efficiencies and microscopy settings, the absolute values from different days of experiments varied. But, the relative differences (ratio) in a value were consistent between NT and TF perturbations. In order to accurately combine data collected from different days, we used the following normalization procedure. First a global mean of NT across all sub-experiments is calculated. Then data from each sub-experiment was normalized through a multiplication constant to bring the mean value of the NT in the sub-experiment to the global NT mean; this multiplication constant is also applied to all the TF perturbations in the same sub-experiment. After the normalization, all the data across different sub-experiments were pooled. For WT data performed on separate days, we combined the data as follows. We matched the pairs of cells with the same PD, and using a scale factor, we scaled the pairs so we minimized the difference between their medians. After the normalization, all the data across different sub-experiments were pooled.

For continuous data, a Wilcoxon rank-sum test was used to compute p values. For nominal data (KI67 and beta-galactosidase positive rates), binomial distribution was used to compute p values. * p values < 0.05, ** p < 0.01, and *** p values < 0.001.

### Gene lists for the cluster heatmaps

To create the cluster heatmaps, a collection of sources was used to generate the gene lists. For proteasome genes, all human proteasome genes were included. For mitochondria and metabolism related genes, all mitochondrial genes (except those encoding tRNA) and the genes derived from the KEGG pathway for “KEGG_CITRATE_CYCLE_TCA_CYCLE”, ID M3985 were included. For the cluster heatmaps on TERT, SV40, and RAS cancer expression, the gene list came from Danielsson et al.^45^. For genes commonly differentially expressed in cancer, the gene list came from Xu et al.^48^. The log 2 fold change for every gene in each sub-list was found for WT passaged cells and TF perturbations, with no p value cut off. Then, fold changes for the genes were clustered using Euclidean distance as the distance metric.

### Detailed list of materials used

| Material name | Manufacturer | Identifier | Concentration/<br>amount used | What used<br>for |
| --- | --- | --- | --- | --- |
| Phospho-Histone H2A.X (Ser139)<br>Rabbit mAb | Cell Signaling<br>Technology | 9718S | 500x | antibody<br>staining |
| Ki-67 Mouse mAb | Cell Signaling<br>Technology | 9449S | 10,000x | antibody<br>staining |
| Histone H3K9me3 (trimethyl Lys9) | Invitrogen | PA5-31910 | 500x | antibody<br>staining |
| Histone H3K27me3 (trimethyl<br>Lys27) Rabbit mAb | Cell Signaling<br>Technology | 9733S | 2,000x | antibody<br>staining |
| Bovine Serum Albumin | Fisher BioReagents | BP9703-100 | 0.5 - 2 % | antibody<br>staining |
| μ-Slide 8 Well chambered slide | ibidi | 80826 | N/A | antibody<br>staining |
| Alexa Fluor™ 488 donkey<br>anti-rabbit antibody | Invitrogen | R37118 | 1 drop per mL<br>antibody dilution<br>buffer | antibody<br>staining |
| Alexa Fluor™ 488 donkey<br>anti-mouse IgG | Invitrogen | A21202 | 2,000x | antibody<br>staining |
| Alexa Fluor™ 546 Phalloidin | Invitrogen | A22283 | 2,000x | antibody staining |
| H3K9me3 Polyclonal Antibody | Invitrogen | PA5-31910 | 500x | antibody staining |
| Triton X-100 | Sigma | X100-100mL | 0.10% | antibody staining |
| 10 % Formaldehyde Solution (w/v)<br>Methanol-free | Thermo Scientific | 28906 | 4% | antibody staining |
| Hoechst 33342 | Thermo Scientific | 62249 | 10,000x | antibody staining |
| XL1-Blue Chemically Competent cells | Macrolab, UC Berkeley | N/A | N/A | Bacterial cloning |
| Hemocytometer, Neubauer Improved | Bulldog Bio | DHC-N01 | N/A | Cell counting |
| Trypan Blue Stain (0.4%) | Gibco | 15250-061 | 2x | Cell counting |
| Primary Dermal Fibroblast Normal;<br>Human,<br>Neonatal (HDFn) | ATCC | PCS-201-010 | N/A | Cell culture |
| Fibroblast growth kit low serum | ATCC | PCS-201-041 | N/A | Cell culture |
| Penicillin-Streptomycin-Amphotericin B Solution | ATCC | PCS-999-002 | 1000x | Cell culture |
| Phenol Red | ATCC | PCS-999-001 | 1000x | Cell culture |
| Fibroblast Basal Medium | ATCC | PCS-201-030 | N/A | Cell culture |
| Fetal Bovine Serum qualified, USA origin | Thermo Fisher Scientific | 26140087 | N/A | Cell culture |
| Minimum Essential Medium | Gibco | 11095-080 | N/A | Cell culture |
| Dulbecco's Modified Eagle Medium | Gibco | 11965-092 | N/A | Cell culture |
| Puromycin | Sigma-Aldrich | 540411 | 2<br>micrograms/mL<br>final | Cell culture |
| Lenti-X™ 293T Cell Line | Takara | 632180 | N/A | Cell culture |
| pJKNp44 dCas9 VPR;<br>pHR-SFFV-HA-NLS-dSpCas9(D10A/H840)-VPR-2PA-BFP | Weissman lab | Link | N/A | CRISPR VPR |
| pMH0001<br>UCOE-SFFV-dCas9-BFP-KRAB | Weissman lab | Link | N/A | CRISPR KRAB |
| pJR89 insert for dual sgRNA plasmid | Weissman lab | Addgene 140096 | N/A | dual sgRNA |
| pJR85 dual sgRNA plasmid | Weissman lab | Addgene | N/A | dual sgRNA |
|  |  | 140095 |  |  |
| QIAamp DNA Blood Mini Kit | Qiagen | 51104 | N/A | gDNA extraction |
| LysoTracker™ Red DND-99 | Invitrogen | L7528 | N/A | lysosome staining |
| TMRE-Mitochondrial Membrane Potential Assay Kit | Abcam | ab113852 | N/A | mitochondria membrane potential stain |
| Promega GloMax plate reader | Promega | N/A | N/A | Proteasome fluorescence assay |
| Proteasome-Glo Chymotrypsin-like cell-based assay | Promega | G8660 | N/A | Proteasome fluorescence assay |
| SuperScript IV Reverse Transcriptase | Invitrogen | 18090050 | N/A | qPCR |
| RNaseOUT Ribonuclease Inhibitor | Invitrogen | 10777-019 | 20x | qPCR |
| KAPA SYBR FAST Universal MasterMix | Roche | KK4602 | 2x | qPCR |
| RNeasy Plus Mini Kit | Qiagen | 74134 | N/A | RNA extraction |
| Chromium Single Cell 3' Kit v2 | 10x Genomics | 120237 | N/A | scRNA-seq library prep |
| Chromium Next GEM Single Cell 5' Kit v1.1 | 10x Genomics | 1000167 | N/A | scRNA-seq library prep |
| Senescence β-Galactosidase Staining Kit | Cell Signaling Technology | 9860 | N/A | Senescence staining |
| pMJ114 (Plasmid #85995); sgRNA, bovine promoter | Weissman lab | Addgene 85995 | N/A | sgRNA backbone |
| pMJ117 (Plasmid #85997); sgRNA human promoter | Weissman lab | Addgene 85997 | N/A | sgRNA backbone |
| pMJ179 (Plasmid #85996) | Weissman lab | Addgene 85996 | N/A | sgRNA backbone |
| TransIT-LT1 transfection reagent | Mirus | MIR 2304 | 15 uL per 6-well; | transfections |

### qPCR primers

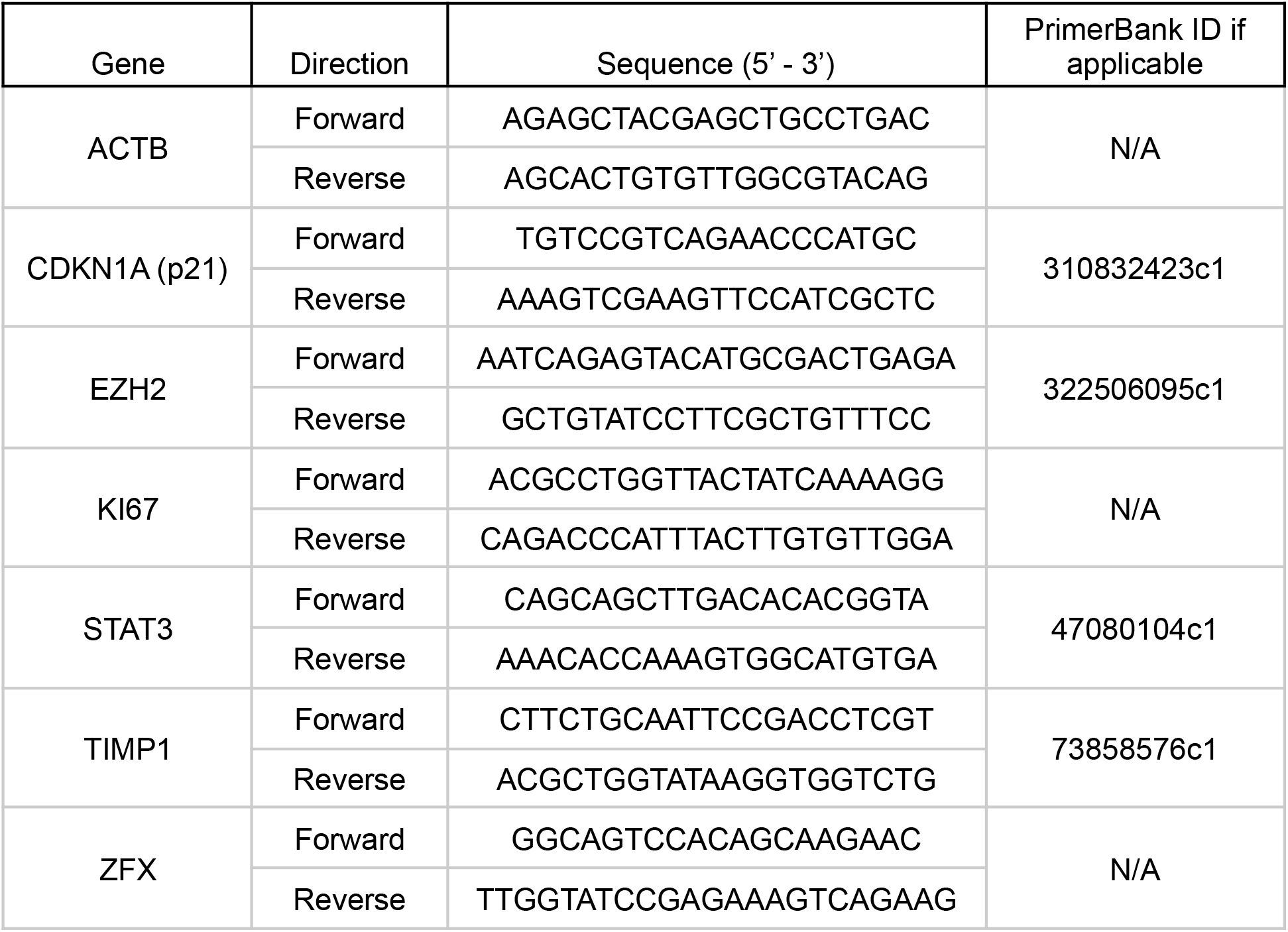

