## Supplemental Figures for "Systematic Approach Identifies Multiple Transcription Factor Perturbations That Rejuvenate Replicatively Aged Human Skin Fibroblasts"

A.

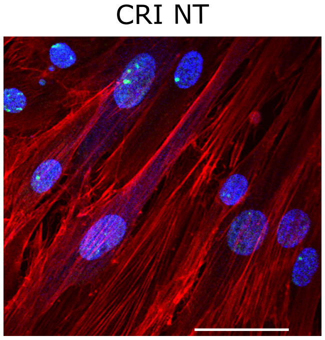


B.

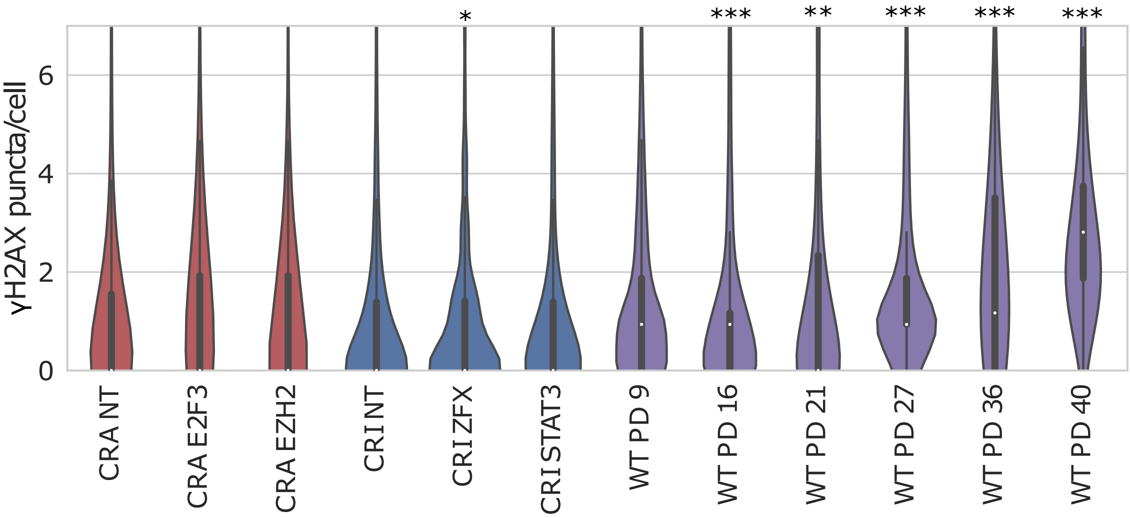


Figure S1. Total γH2AX DNA foci do not change considerably in TF perturbations, but there is an increase in late passage WT cells.

A. CRI NT γH2AX DNA foci imaging. Blue is Hoechst staining DNA, red is a phalloidin staining actin, and green are γH2AX DNA foci, 50 uM scale bar.

B. Quantification of γH2AX puncta per cell; N > 500 cells per TF perturbation, N > 240 cells for each WT PD. Significance was calculated by a Wilcoxon rank-sum test, comparing TF to NT for CRA and CRI, and later passages to PD 9 for WT. *p < 0.05, **p < 0.01, ***p < 0.001.

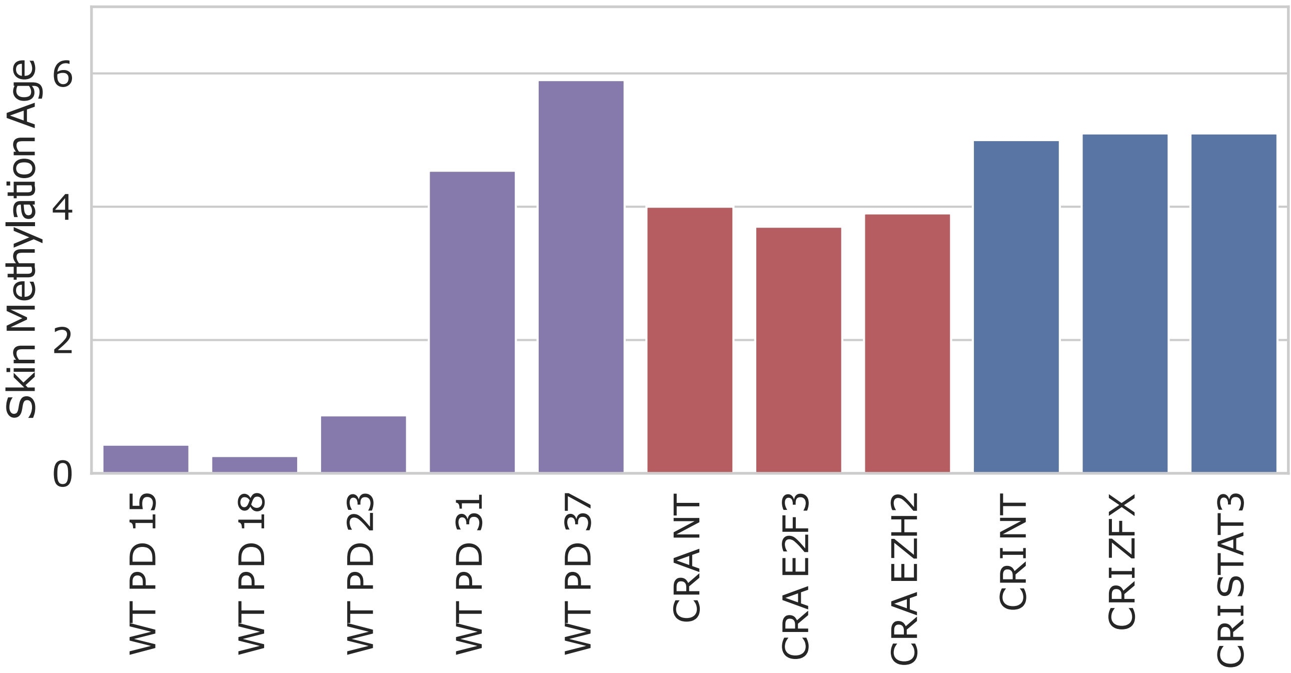


Figure S2. Skin methylation clock analysis of WT passaged cells, CRA TF perturbations, and CRI TF perturbations.
WT passaged cells have a progressively higher (older) methylation age. None of the TF perturbations have altered methylation ages compared to NT control cells.

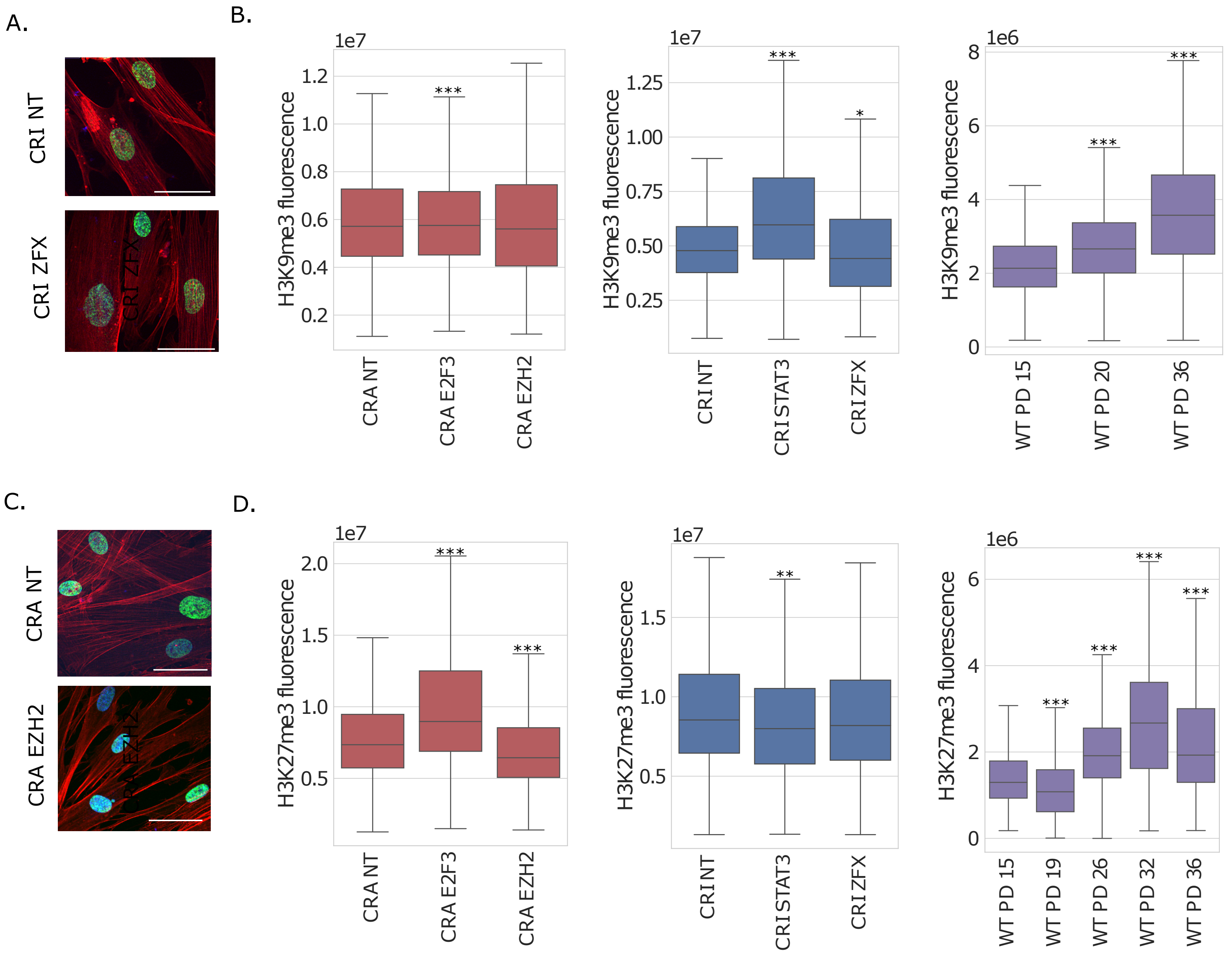


Figure S3. Total expression of epigenetic markers H3K9me3 and H3K27me3 are variable across TF perturbations but trend upwards in later WT passages.

A. CRI NT and CRI ZFX H3K9me3 staining; blue is Hoechst, red is a phalloidin staining actin, and green is H3K9me3, 50 uM scale bar.

B. Quantification of H3K9me3 total fluorescence per nucleus. For CRA and CRI cells, N > 250 cells per sgRNA and N > 1,170 cells per WT PD; significance was calculated by a Wilcoxon rank-sum test compared to NT for CRA and CRI, and PD 15 for WT. * $p < 0.05$, ** $p < 0.01$, *** $p < 0.001$.

C. CRA NT and CRA EZH2 H3K27me3 staining; blue is Hoechst, red is phalloidin staining actin, and green is H3K27me3, 50 uM scale bar.

D. Quantification of H3K27me3 total fluorescence per nucleus. For CRA and CRI cells, N > 250 cells per sgRNA and N > 600 cells per WT PD; significance was calculated by a Wilcoxon rank-sum test compared to NT for CRA and CRI, and PD 15 for WT. * $p < 0.05$, ** $p < 0.01$, *** $p < 0.001$.

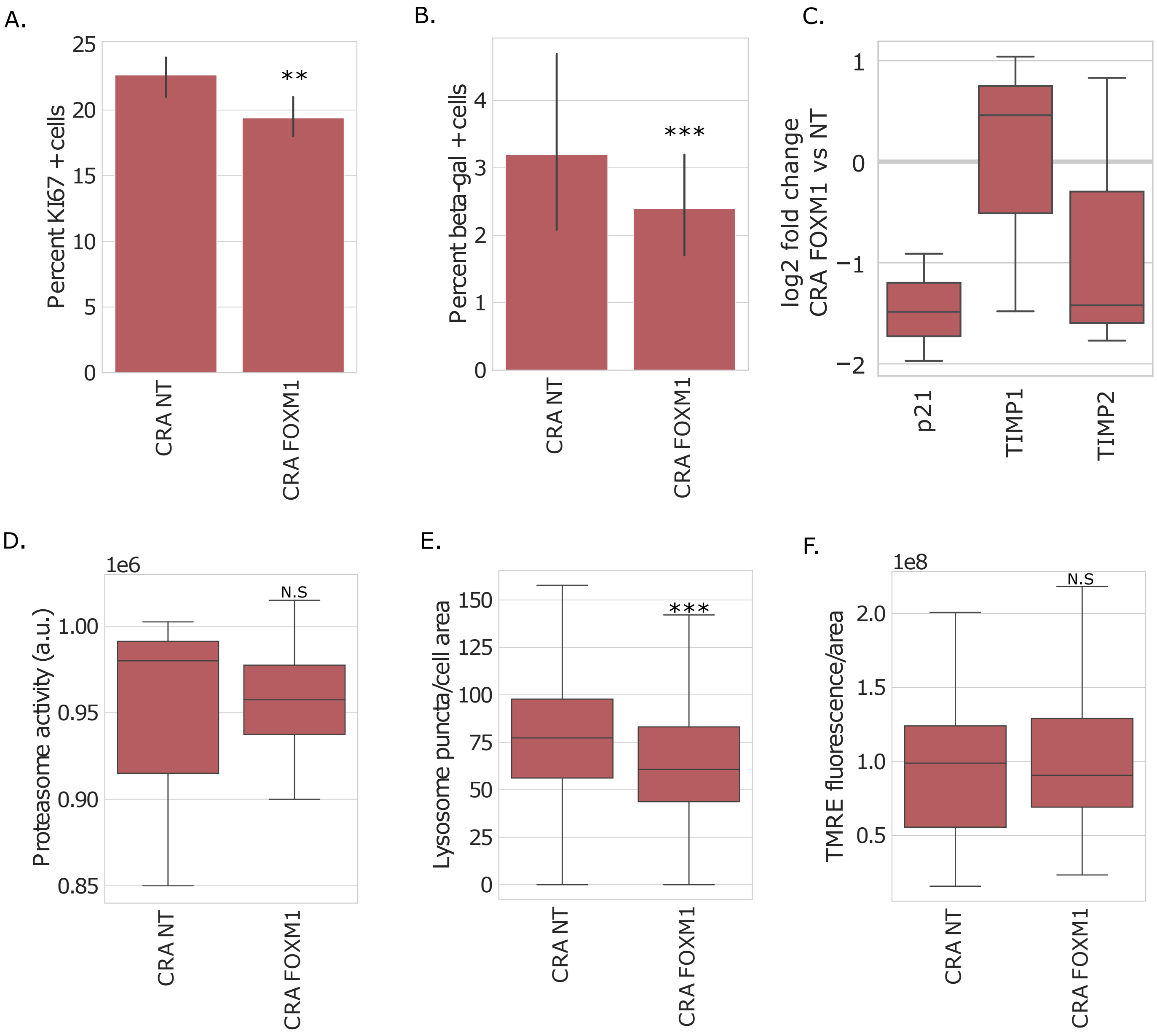


Figure S4. CRA FOXM1 late passage cells phenocopy some passaged cell rejuvenation phenotypes, but not all.
A. Percent KI67 positive cells.
B. Percent beta-gal positive cells.
C. Quantitative PCR for common senescence genes, with the log two fold change (log2fc) relative to CRA NT.
D. Proteasome activity.
E. Lysosome puncta per cell area, as measured with LysoTracker Red.
F. TMRE fluorescence per cell area.
For continuous variables, significance was calculated by Wilcoxon rank-sum test. For binary outcomes, significance was calculated based on binomial distribution. *p $<$ 0.05, **p $<$ 0.01, ***p $<$ 0.001.
